# A Geometric Framework for Understanding Dynamic Information Integration in Context-Dependent Computation

**DOI:** 10.1101/2021.02.09.430498

**Authors:** Xiaohan Zhang, Shenquan Liu, Zhe Sage Chen

## Abstract

Prefrontal cortex plays a prominent role in performing flexible cognitive functions and working memory, yet the underlying computational principle remains poorly understood. Here we trained a rate-based recurrent neural network (RNN) to explore how the context rules are encoded, maintained across seconds-long mnemonic delay, and subsequently used in a context-dependent decision-making task. The trained networks emerged key experimentally observed features in the prefrontal cortex (PFC) of rodent and monkey experiments, such as mixed-selectivity, sparse representations, neuronal sequential activity and rotation dynamics. To uncover the high-dimensional neural dynamical system, we further proposed a geometric framework to quantify and visualize population coding and sensory integration in a temporally-defined manner. We employed dynamic epoch-wise principal component analysis (PCA) to define multiple task-specific subspaces and task-related axes, and computed the angles between task-related axes and these subspaces. In low-dimensional neural representations, the trained RNN first encoded the context cues in a cue-specific subspace, and then maintained the cue information with a stable low-activity state persisting during the delay epoch, and further formed line attractors for sensor integration through low-dimensional neural trajectories to guide decision making. We demonstrated via intensive computer simulations that the geometric manifolds encoding the context information were robust to varying degrees of weight perturbation in both space and time. Overall, our analysis framework provides clear geometric interpretations and quantification of information coding, maintenance and integration, yielding new insight into the computational mechanisms of context-dependent computation.

## INTRODUCTION

Cognitive flexibility is an important characteristic that enable animals or humans to selectively switch between sensory inputs to generate appropriate behavioral responses (Diamond, 2013; Scott, 1962; Miyake et al., 2012). This important process has been associated with various goal-directed behaviors, including multi-tasking, decision-making (Thea, 2012; Dajani and Uddin, 2015; Le et al., 2018; Pezzulo et al., 2014). Impaired cognitive flexibility has been observed among individuals with mental illness, such as schizophrenia (Woodward et al., 2012; Maud et al., 2012) and those at risk for mental disorder (Murphy et al., 2012; Chamberlain et al., 2007; Vaghi et al., 2017). Therefore, identifying the computational principle underlying cognitive flexibility may improve our understanding of brain dysfunction. The prefrontal cortex (PFC) is known to contribute to cognitive flexibility, serving as the main storage of temporary working memory (WM) to represent and maintain contextual information (Baddeley, 2003; Miller, 2000; Todd et al., 2009). Neurophysiological recordings have shown that single PFC cells respond selectively to different task-related parameters (White and Wise, 1999; Eiselt and Nieder, 2016; Hyman et al., 2013; Machens et al., 2010; Rigotti et al., 2013), and the activity of PFC pyramidal neurons can maintain WM to perform context-dependent computation (Wallis et al., 2001). However, due to the heterogeneity and diversity of single-neuron responses, it remains challenging to understand how task-modulated single-neuron activities integrate task-related information to guide subsequent decision making. To address this knowledge gap, researchers relied on population coding to understand the maintenance and manipulation of context information in decision-making tasks (Meyers et al., 2008; Cichy et al., 2014; King and Dehaene, 2014; Lundqvist et al., 2016). In the literature, several computational theories and analysis methods have been proposed (Wu et al., 2020; Mante et al., 2013). However, how the sensory information is integrated via dynamic population coding in a context-dependent manner remains poorly understood. Meanwhile, an intuitive and interpretable dynamical systems framework for context-dependent WM and decision-making is lacking.

Recurrent neural networks (RNNs) have been widely used for modeling a wide range of neural circuits, such as the PFC and parietal cortex, in performing various cognitive tasks (Rajan et al., 2016; Hennequin et al., 2014; Sussillo et al., 2015). However, computational mechanisms of RNNs in performing those tasks remain elusive because of the black-box modeling. In this paper, we trained a RNN to perform a delayed context-dependent task (Figure 1*A*) and proposed a geometric analysis framework to understand dynamic population coding and information integration. We found that the trained RNN captured critical physiological properties consistent with reported experimental data. Additionally, the trained RNN showed some emergent features of neuronal activity observed in the PFC, such as the mixed-selectivity, sparse representation, and sequential activity. Based on dimensionality reduction of population responses, we defined task epoch-specific subspaces and dynamic attractors during the sensory integration epoch, and showed that context-configured network state is temporally tuned to regulate sensory integration to guide decision-making. Together, our analysis framework not only helps uncover the computational mechanisms of encoding and maintenance of context information in decision-making, and but also helps illustrate information integration based on interpretable geometric concepts.

**Figure 1:**
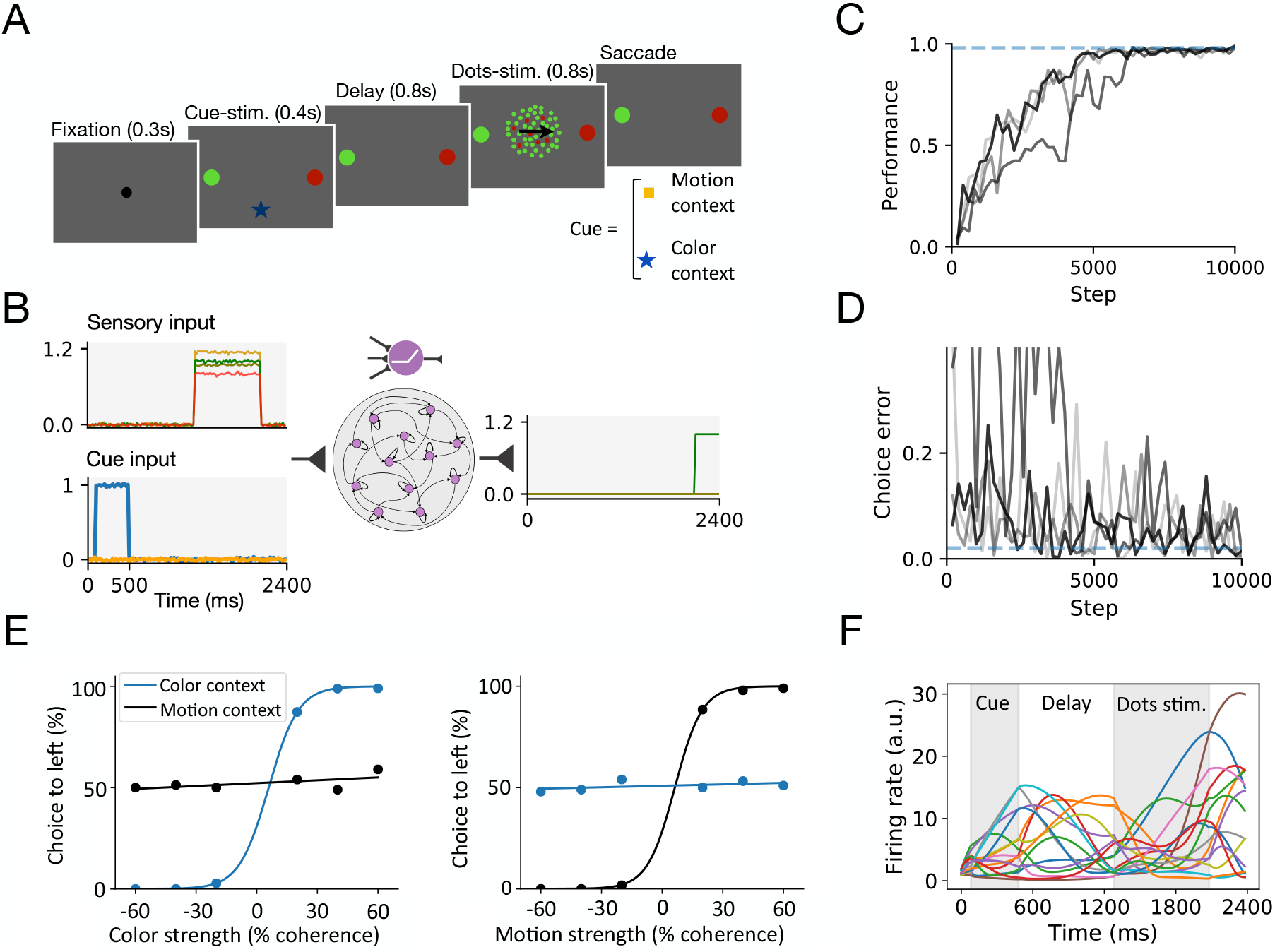
Trained RNN to Perform the Delayed Context-dependent Integration Task. **(A)** Behavioral task description. Monkey was trained to discriminate, depending on the contextual cue, either the predominant color or predominant motion direction of randomly-moving dots, and to further indicate its decision with a saccadic eye movement to a choice target. The cue stimulus onset determined the current context, which was characterized by different shapes and colors of the fixation point. The cue stimulus was followed a fixed-delay epoch, and then followed by randomly-moving dots stimuli. The monkey was rewarded for a saccade to the target matching the current context. **(B)** Schematic of a fully connected, nonlinear RNN in context-dependent computation. The network received time-varying sensory and cue inputs, and produced a desired output. The two input channels encoded sensory information and context cues, and the two output channels encoded the response direction. **(C-D)** RNN learning curve (**C**) and the performance curve (**D**). Training was completed once both quantities reached the convergence criterion (blue horizontal dashed lines). **(E)** Psychometric curves in a delayed context-dependent integration task. The probability of a correct direction judgment is plotted as a function of color (*Left*) and motion (*Right*) coherence in color-context (blue) and motion-context (black) trials. (**F**) The activities of representative units indicated by different colors. The first grey shading area indicates cue stimulus epoch and the second shading area indicates the presentation of random dots (*i.e.*, integration of sensory stimulus epoch).

## RESULTS

### Trained RNN for Performing a Delayed Context-Dependent Integration Task

We trained the RNN to perform a delayed context-dependent WM or decision making task (Figure 1*B*). At each trial, the network received two types of noisy inputs: sensory stimulus and cue stimulus. The sensory input units encoded the momentary motion and color evidence toward two target directions. The cue input units encoded the contextual signal, instructing the network to discriminate specific type of sensory input. The choice output units encoded the response direction. All units had non-negative and non-saturating firing rates to mimic the properties of biological neurons (Priebe and Ferster, 2008; Abbott and Chance, 2005).

Upon successful convergence of RNN training (Figures 1*C* and 1*D*), psychometric tests showed that the trained RNN captured critical physiological properties consistent with experimental findings (Figure 1*E*). For example, the trained network achieved better performance with higher color coherence stimulus in the color context, but not in the motion context; and vice versa (Figure 1*E*). Units in the trained RNN showed diverse firing rate profiles at different task epochs (Figure 1*F*). Furthermore, we analyzed the impact of the proportion of zero recurrent weights on the task performance (Figure 1*A*). The trained RNN exhibited a strong self-connection (Figure 1*B*). Weight perturbation analyses were also used to assess the stability of the trained RNN (Figure S2).

### Single Unit Responses

#### Mixed-selectivity

Mixed selectivity of PFC neurons is important for implementing complex cognitive functions, manifesting itself as an ‘adaptive coding’ strategy (Arnal et al., 1998). We found that many units of the trained RNN exhibited mixed selectivity for task-related variables (Figure 2*A*). A unit was said to be selective to a task-related variable if it responded differently to the values of the parameters characterizing that variable. We classified the units based on their responses to different task parameters, and found four distinct types of mixed-selective units from the trained RNN. (i) Some units exhibited mixed selectivity to task rules (both color context cue and motion context cue), such as unit 3. (ii) Some units exhibited mixed selectivity to both color sensory stimuli (red and green), such as unit 8. (iii) Some units exhibited mixed selectivity to both directions of coherent motion, such as unit 12. (iv) Some units exhibited selectivity to both task rules and sensory stimuli. For example, unit 5 and unit 9 responded to both the color cue stimuli and color sensory stimuli. Unit 4 and unit 13 responded to both the motion cue stimuli and motion sensory stimuli.

**Figure 2:**
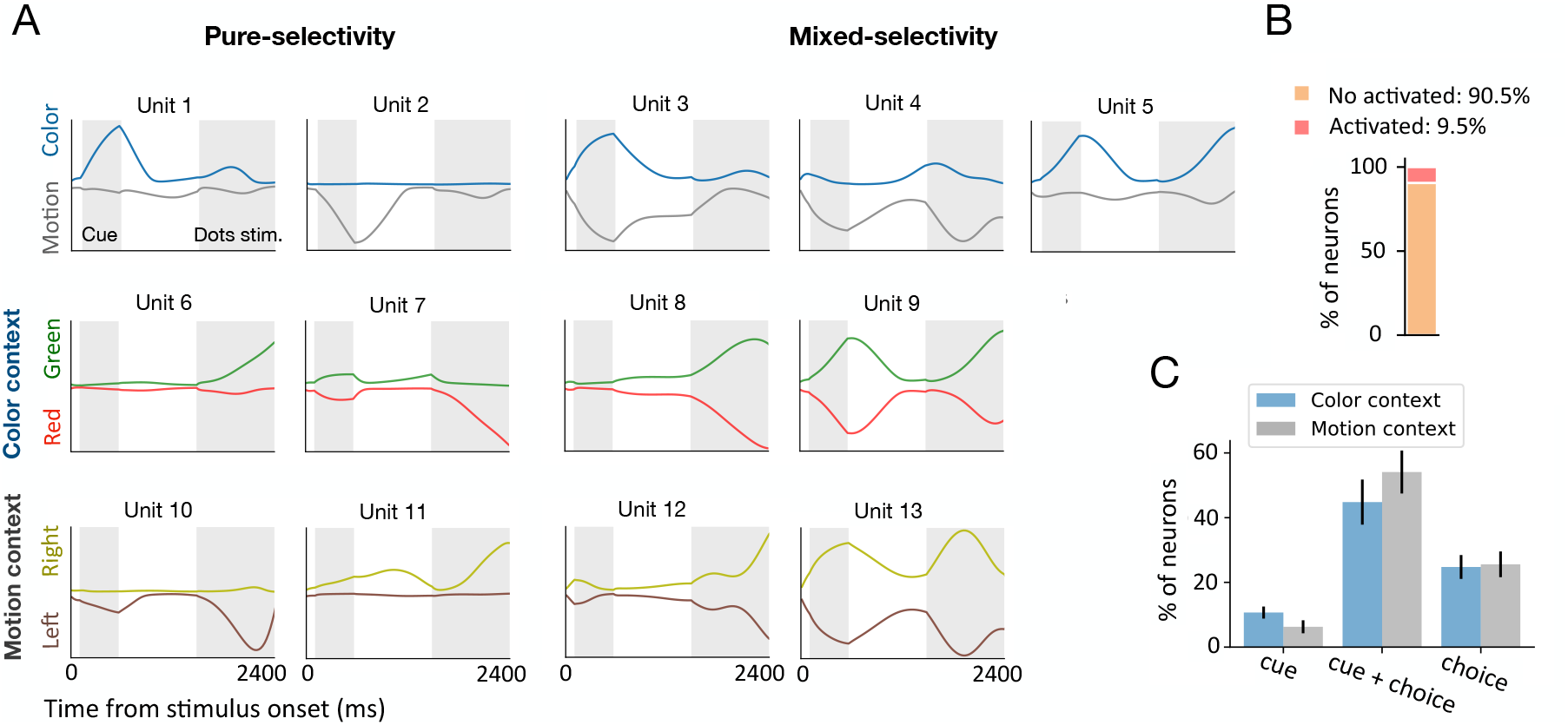
Single Unit Responses. **(A)** Single unit responses under different task conditions, as indicated by different colors. Two shaded areas represent the period of contextual cue stimulus presentation and sensory stimulus presentation, respectively. *Left panel* : Responses of units showed pure-selectivity to the task-related variable. Unit 1 preferred the color cue, unit 2 preferred the motion cue, unit 6 preferred green, unit 7 preferred red, unit 10 preferred the leftward direction, and unit 11 preferred the rightward direction. *Right panel* : Different task-related variables were mixed. Unit 3 showed mixed selectivity for the color cue and motion cue. Units 4 and unit 5 showed mixed selectivity for both the context cue and sensory stimulus. Unit 8 showed mixed selectivity for two task variables (green and red) for the color context. Unit 12 showed mixed selectivity for two task variables (left and right) for the motion context. Unit 9 showed mixed selectivity for three task features, such as the color context cue, green, and red sensory stimulus. Similarly, unit 13 showed mixed selectivity for the motion context cue, leftward direction, and rightward direction. **(B)**The percentage of units in the trained RNN that were activated to perform a delay context-dependent decision making task, averaging over 20 training configurations. **(C)** For a given context, the percentage of units that were selective to the context cue, sensory input, and their combinations. Error bar indicates SEM over 20 different training configurations.

Although single-unit activity could be tuned to mixtures of multiple task-related variables, some other units were modulated primarily by only one of the task variables (what we will call ‘pure-selectivity’ units). For example, unit 1 was primarily selective to the color context cue, and unit 2 was selective to the motor cue. Some units only responded to sensory stimuli, such as unit 6, unit 7, unit 10, and unit 11 (Figure 2*A*). In the trained RNN, only a small subset of units showed activations to task variables (Figure 2*B*). Among those activated units, the number of units encoding the choice was much more than that of units encoding the context cue (Figure 2*C*). One possible explanation is that it was more difficult to integrate noisy sensory information than to distinguish the context information, so more units were recruited to process sensory information.

We have mainly introduced four types of mixed selectivity units, with varying degrees of selective responses to different task-related variables. However, the diversity of mixed selectivity responses often resulted in response properties that were not easily interpretable. For example, in the color context, unit 9 simultaneously responded to the color cue stimulus and green signal, which also responded to the red signal. Specifically, units with such mixed selectivity behaved the same way in the same context, causing a difficulty of interpretation. This suggests that the activity of individual mixed-selectivity units could not fully disambiguate the information; only when pooling information from multiple units, the ambiguity of information encoded by mixed-selective units can be eliminated—supporting the necessity of population coding by ensembles of neurons (Rafael, 2015).

#### Sparse representation

On average, only 9.5% recurrent units in the trained RNN were activated to perform the task (Figure 2*B*), indicating the notion of sparse representations or sparse coding (Olshausen and Field, 2004). We found that mixed-selectivity units accounted for approximately 50% of the total activated units (Figure 2*C*), and the information represented by mixed-selectivity units was highly correlated. This required the network to decorrelate the representations sufficiently and thereby increase the classification capacity. Additionally, sparse coding formed an easy-to-interpret representation of input combinations, ensuring the trained network to discriminate between similar inputs. Notably, the level of sparsity in our network was close to an optimal value that minimizes the classification error (Barak et al., 2013a).

### Population Response

We further studied the neural representation at the population level. The dynamics of population activity can be characterized through the high-dimensional state space **x**(*t*) ∈ ℝ^*N*^. The time-varying population activity can be visualized as a trajectory within the lower-dimensional subspace, and the distance between the points in the subspace reflects the population response difference.

#### Cue Processing

To examine cue processing, we reported the dynamics of population activity throughout the cue stimulation and delay epochs. We performed principal component analysis (PCA) to identify a three-dimensional neural subspace, which was spanned by the first three principal component (PCs) that accounted for about 92% of neural activity variance (Figure 3*A*). During the cue presentation epoch, the network started at the the same subspace and then evolved along different trajectories based on the contextual cues (Figure 3*B*). Further, the distance between two state trajectories increased (Figure 3*C*, blue curve), indicating that these two trajectories were continuously divergent during the cue stimulus epoch. Moreover, we defined the “energy” of the population activation state using the average activity of the total number of activated units (Figure 3*C*, grey curve).

**Figure 3:**
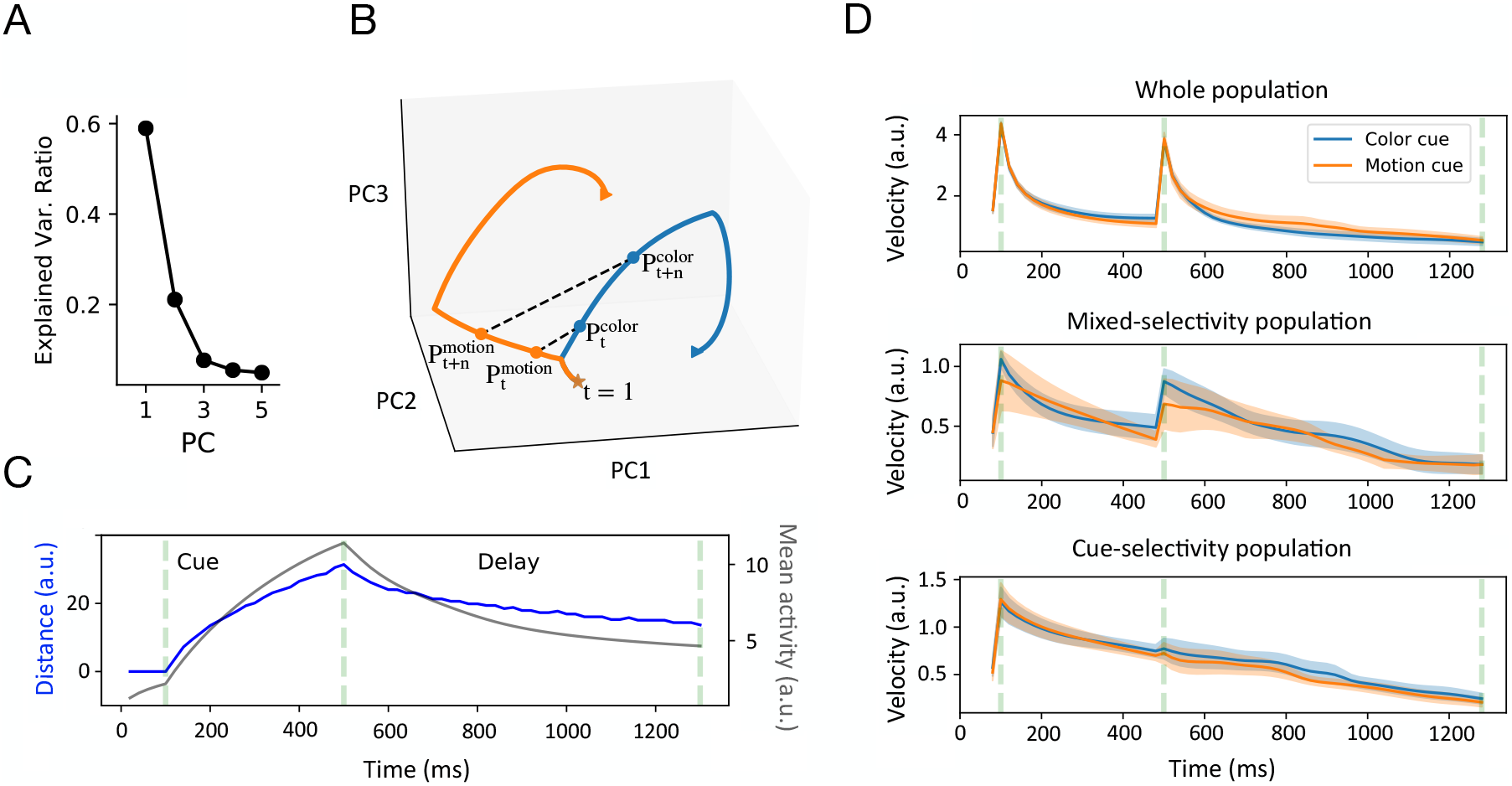
Neural Population Dynamics. **(A)** Ratio of explained variance of the first five PCs of neural subspace during the cue stimulus and delay epochs. **(B)** Two different neural trajectories in a three-dimensional subspace during the cue stimulus and delay epochs. Orange and blue curves correspond to the motion and color contexts, respectively. The three-dimensional distance between two context-specific states at time *t* characterized the similarity of their population responses: 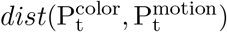. In a context-specific state trajectory, we calculated the distance between states at time *t* and *t* + *n*, quantifying the change in trajectory position as a function of time, or velocity: 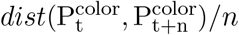. **(C)** Distance between two context-specific trajectories in the cue stimulus and delay epochs as a function of time (blue curve). For comparison, the overall mean network activity (or network energy) is shown in gray curve (right axis). **(D)** The top panel plots the velocity of the temporal evolution of overall population state (*i.e.*, 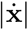) under color context (blue) and motion context (orange). Shaded area denotes SEM over 20 training configurations. The middle and bottom panels show the state velocity evolution of mixed-selectivity population and cue-selectivity population, respectively.

We found that the distance curves between two cue-specific trajectories was temporally aligned with the increase of energy level during the cue stimulus and delay epochs. During the delay epoch, the network settled in a low-energy state. Moreover, the distance between two trajectories reached a peak value at the end of the cue stimulus presentation, then decreased during the delay epoch until reaching a plateau. However, the plateau value was greater than zero (a similar level as at time *t* = 200 ms), suggesting that the difference in cue-specific trajectories still existed to to distinguish the context conditions.

We computed the “velocity” of population activity as a function of time. Before the cue stimulus appeared, the overall population activity had a rapid acceleration to reach a peak (Figure 3*D*, top panel), and then the velocity of population activity gradually decayed to the pre-stimulus baseline level, suggesting that cue-specific trajectories were separated at a stable velocity in the late stage of cue-stimulus. During the delay epoch, the velocity of population activity jumped to a large value again and then dropped rapidly. The phenomenon was primarily contributed by mixed-selective units. Specifically, we plotted the respective velocities of population activity with pure-selectivity and mixed-selectivity for comparison (Figure 3*D*). We observed that the velocity of the population that was only sensitive to the context cue decreased continuously throughout the cue stimulus and delay epochs. In contrast, for the mixed-selectivity population, there was a jump point in the velocity curve at the beginning of the delay epoch. Therefore, the velocity is sensitive to change of epoch-wise population activity, and provides an informative measure of the population dynamics. Motivated by the local optogenetic manipulation in animal experiments (Gray et al., 2017), we conducted temporally local weight perturbation analyses during the delay epoch, and found that the RNN performance was more robust with local perturbation (Figure S3) than with global perturbation (Figure S2).

#### Sequential activity

Neural sequences are emergent properties of RNNs in many cognitive tasks, which has been thought to be a common feature of population activity during a wide range of behaviors (Fiete et al., 2010; Rajan et al., 2016; Orhan and Ma, 2019). We found that our trained RNN generated emergent sequential activity during the delay epoch (Figure 4*F*). To quantify the sequentiality of neural activity, we calculated the sequentiality index (SI). By virtue of random sampling and Monte Carlo statistics, we found that the trained network has a higher SI (*p* < 0.017, two-sample Kolmogorov-smirnov test) than the untrained network (Figures 4*A* and 4*B*), suggesting that stronger sequential activity emerged from a trained network (Figure 4*F*).

**Figure 4:**
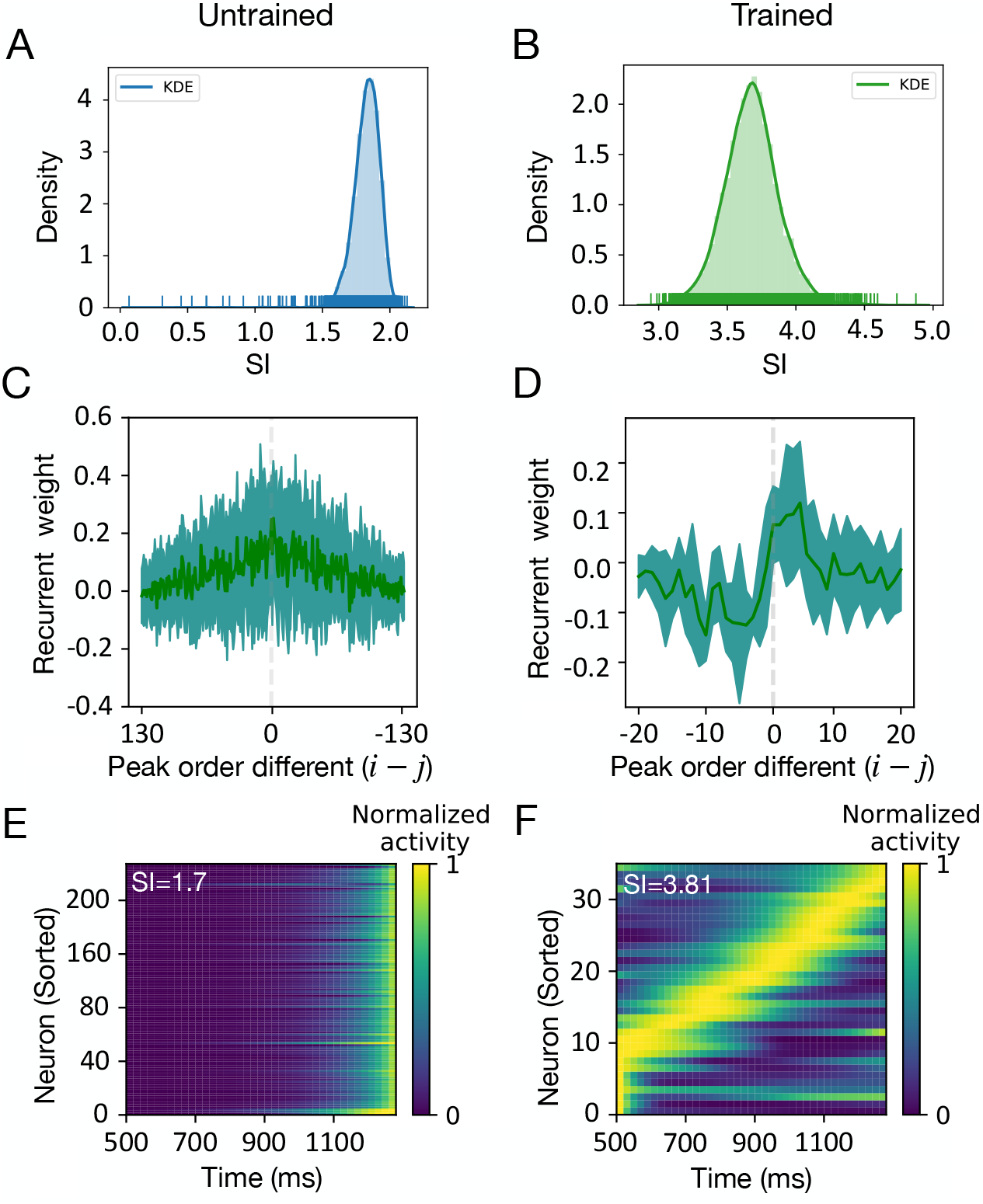
Sequential Neural Representation. **(A-B)** The Monte Carlo distribution of sequentiality index (SI), which was computed by 10000 samples from repeated random sampling. SI was computed from the untrained **(A)** and trained **(B)** networks. **(C-D)** Units were sorted according to their peak time in untrained (**C**) and trained (**D**) networks. The recurrent weights (*W*_*ij*_) were plotted as a function of the peak order difference between pre- and post-synaptic units during the cue-delay epoch. A positive (*i* − *j*) value represents a connection from a pre-to a post-synaptic unit. A negative (*i* − *j*) value represents a connection from a post-to a pre-synaptic unit. Both mean (solid lines) and standard deviation (shaded regions) statistics were computed from multiple trained networks. **(E-F)**Heat maps of normalized unit activity normalized by the peak response (per row) and sorted by the peak time. The activity did not appear ordered in an untrained network **(E)**, whereas sequential activity emerged in the trained network **(F)**.

Next, we investigated the computational mechanism that produces neural sequential activity. Similar to Rajan et al. (2016), we ordered the peak firing time of recurrent units and computed the mean and standard deviations of the recurrent weights 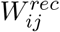. These mean statistic was plotted as a function of the order difference (*i* − *j*) between the *i*-th and *j*-th units (Figures 4*C* and 4*D*). Interestingly, the connection weight of trained RNN showed an asymmetric peak, that is, the connection in ‘forward’ direction (*i.e.*, from earlier-peaking to later-peaking units) was strengthened more than those in ‘backward’ direction (*i.e.*, from later-peaking to earlier-peaking units) (Figure 4*D*). Therefore, this asymmetrical peak weakened the connections between temporally distant units, while strengthening the connections between temporally close units. However, this asymmetric structure was absent in the untrained network (Figure 4*C*), resulting in the loss of sequential activation structure (Figure 4*E* and Figure S4). Put together, the asymmetric weight profile in the trained RNN could prolong responses in later-peaking units, producing the emergent sequential activity.

#### Rotation dynamics

The jPCA is a dimensionality reduction technique that finds an oscillatory structure in the data (Churchland et al., 2012). We performed jPCA on population responses in different contexts and visualized the two-dimensional projections of responses in the jPCA space (Figure 5). Interestingly, the trained RNN showed a strong rotation dynamic similar to those observed in the brain (Sussillo et al., 2015), which is consistent with the findings of (Lebedev et al., 2019).

**Figure 5:**
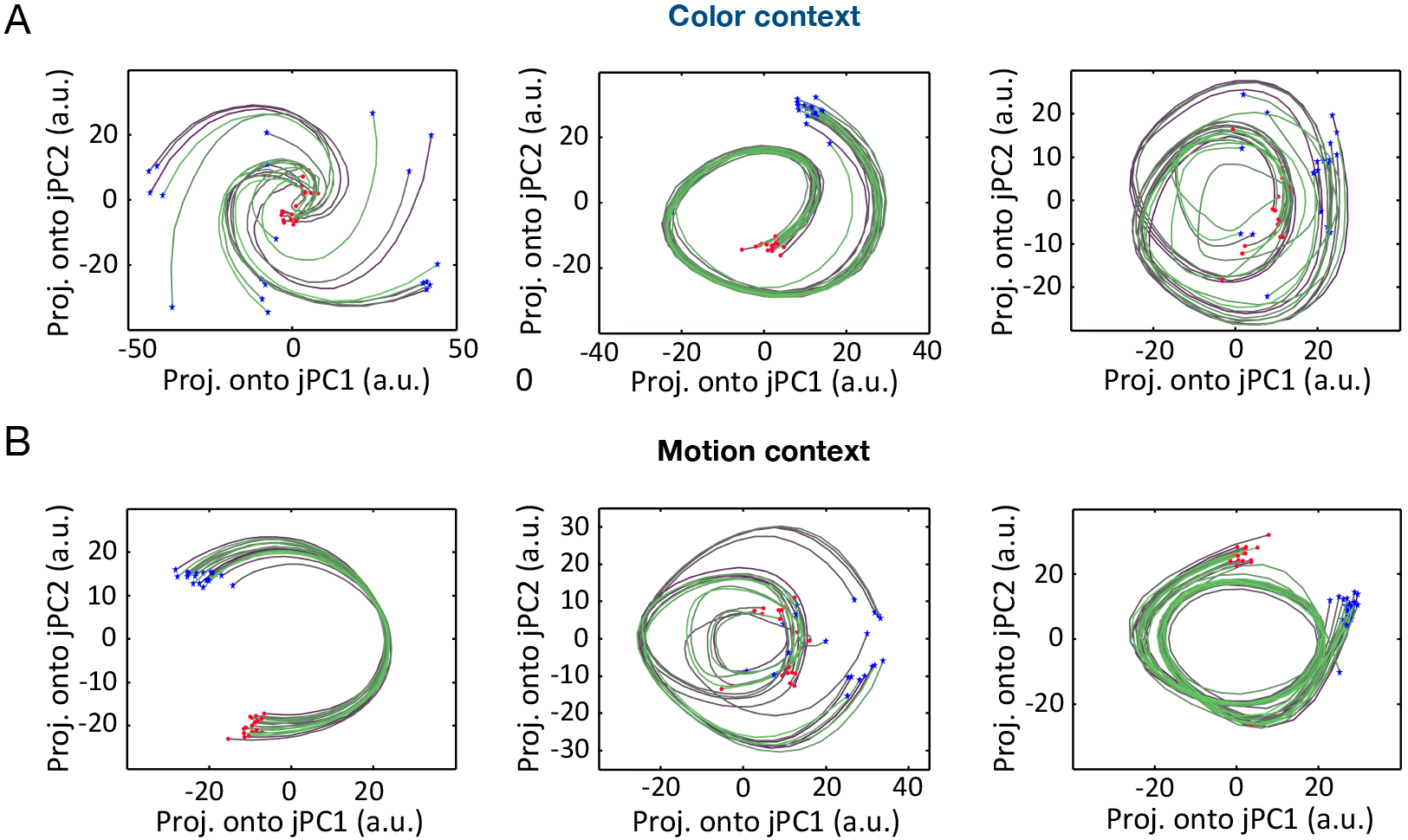
jPCA Projections of the Population Response during the Delay Epoch. **(A)** Three examples of two-dimensional population rotational dynamics in the color context, which corresponded to different training configurations. Each trace represents one computer simulation trial (blue star: trial start point; red circles: trial end point). **(B)** Three examples of two-dimensional population rotational dynamics in the motion context.

### Definition of Task Epoch-specific Subspaces and Axes

To probe how neural population activity dynamically encoded task-related variables, we analyzed the population responses during six different task periods, including four single task epochs (cue stimulus epoch, delay epoch, integration of sensory stimulus epoch, and response epoch) and two cross-epoch periods (Figure 6). As expected, the correlation between population firing during these six periods varied (Figure S5).

**Figure 6:**
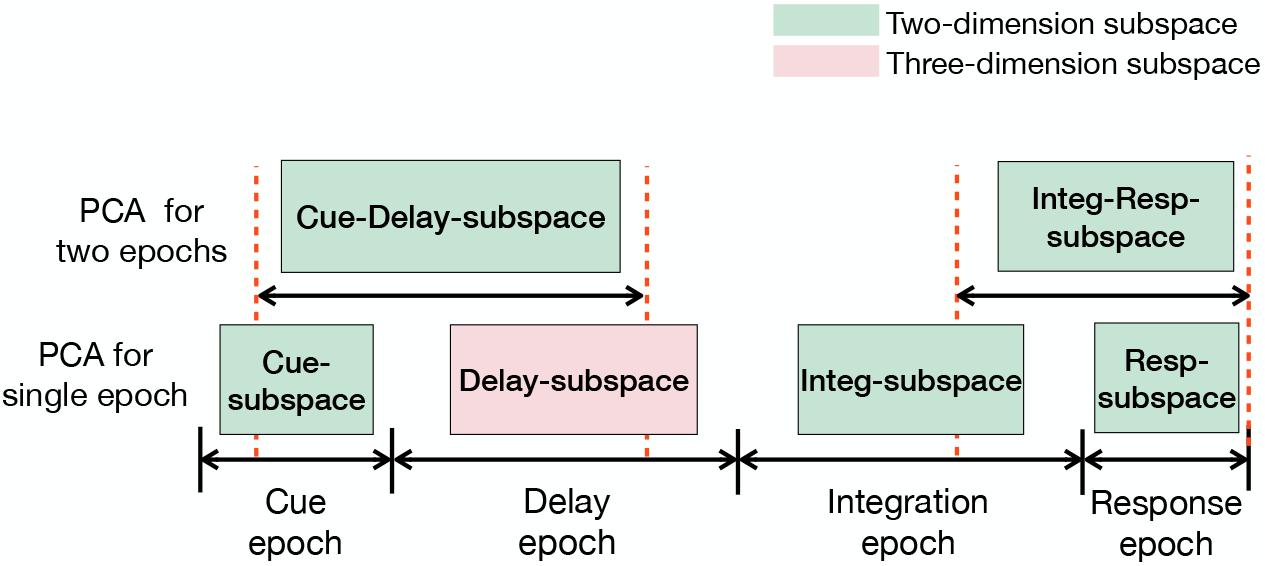
Illustration of Distinct Subspaces Generated in Different Task Epochs. Six subspaces are obtained by performing epoch-wise PCA on population response activities during six different periods, including four single task epochs (Cue epoch, Delay epoch, Integration epoch, and Response epoch) and two cross-epochs (Cue-Delay epoch, and Integration-Response epoch).

We first performed epoch-wise PCA on the population response at single task epochs to generate the corresponding state subspaces (*i.e.*, Cue-subspace, Delay-subspace, Integ-subspace and Resp-subspace). To characterize the evolution of trajectory, we further defined four task-related axes (Table 1): the axis of color context cue (`C-cue-axis`), the axis of motion context cue (`M-cue-axis`), the axis of color choice (`C-choice-axis`), the axis of motion choice (`M-choice-axis`). The definitions of these task epoch-specific subspaces and axes provide a geometric framework for population response analyses. Next, we projected these four task-related axes onto the corresponding state subspaces and examined the neural trajectory in a subspace-specific manner.

**Table 1:** The definition of geometry concepts in five neural subspaces. The matrix **X** varies according to different subspace.

| Figure | Subspace<br>(in two dimensions) | Geometric<br>notion | Definition and interpretation |
| --- | --- | --- | --- |
| Figure 7A | Cue-subspace | C-cue-axis | PCA on matrix $\mathbf{X}_{cue,color}$, the first PC is defined as the C-cue-axis |
| | | M-cue-axis | PCA on matrix $\mathbf{X}_{cue,motion}$, the first PC is defined as the M-cue-axis |
|  |  | cue-PC1<br>cue-PC2 | Performing PCA on population responses during the cue stimulus epoch.<br>The first three PCs are: cue-PC1, cue-PC2, cue-PC3 |
| Figure 7C | Cue-Delay-subspace | cue-delay-PC1<br>cue-delay-PC2 | PCA on population responses across two task epochs (Figure 6). The first three PCs are: cue-delay-PC1, cue-delay-PC2, cue-delay-PC3 |
| Figure 8D | Integ-subspace | C-choice-axis | PCA on matrix $\mathbf{X}_{integ,color}$, the first PC is defined as the C-choice-axis |
| | | M-choice-axis | PCA on matrix $\mathbf{X}_{integ,motion}$, the first PC is defined as the M-choice-axis |
|  |  | integ-PC2<br>integ-PC3 | PCA during the sensory stimulus epoch. The first three PCs are: integ-PC1, integ-PC2, integ-PC3. |
| Figure 9B | Resp-subspace | resp-PC2<br>resp-PC3 | PCA on population responses during the response epoch (Figure 6). The first three PCs are: resp-PC1, resp-PC2, resp-PC3 |
| Figure 9D | Integ-Resp-subspace | integ-resp-PC2<br>integ-resp-PC3 | PCA on population responses across two task epochs (Figure 6). The first three PCs are: integ-resp-PC1, integ-resp-PC2, integ-resp-PC3 |

First, we performed PCA on the population response during the cue stimulus epoch, and the first two principal components (cue-PCs) explained 93.1% of data variance (Figure S6*A*). This indicates that trajectories in the two-dimensional subspace (denoted as Cue-subspace spanned by cue-PC1 and cue-PC2) captured the majority of variance of population responses. We had a similar finding in the Cue-subspace as shown in Figure 3*B*: The network state started from the same location and then produced different trajectories according to different cue stimulus conditions (Figure 7*A*). For a given context, the neural state evolved along a stereotypical trajectory across all sensory stimuli.

**Figure 7:**
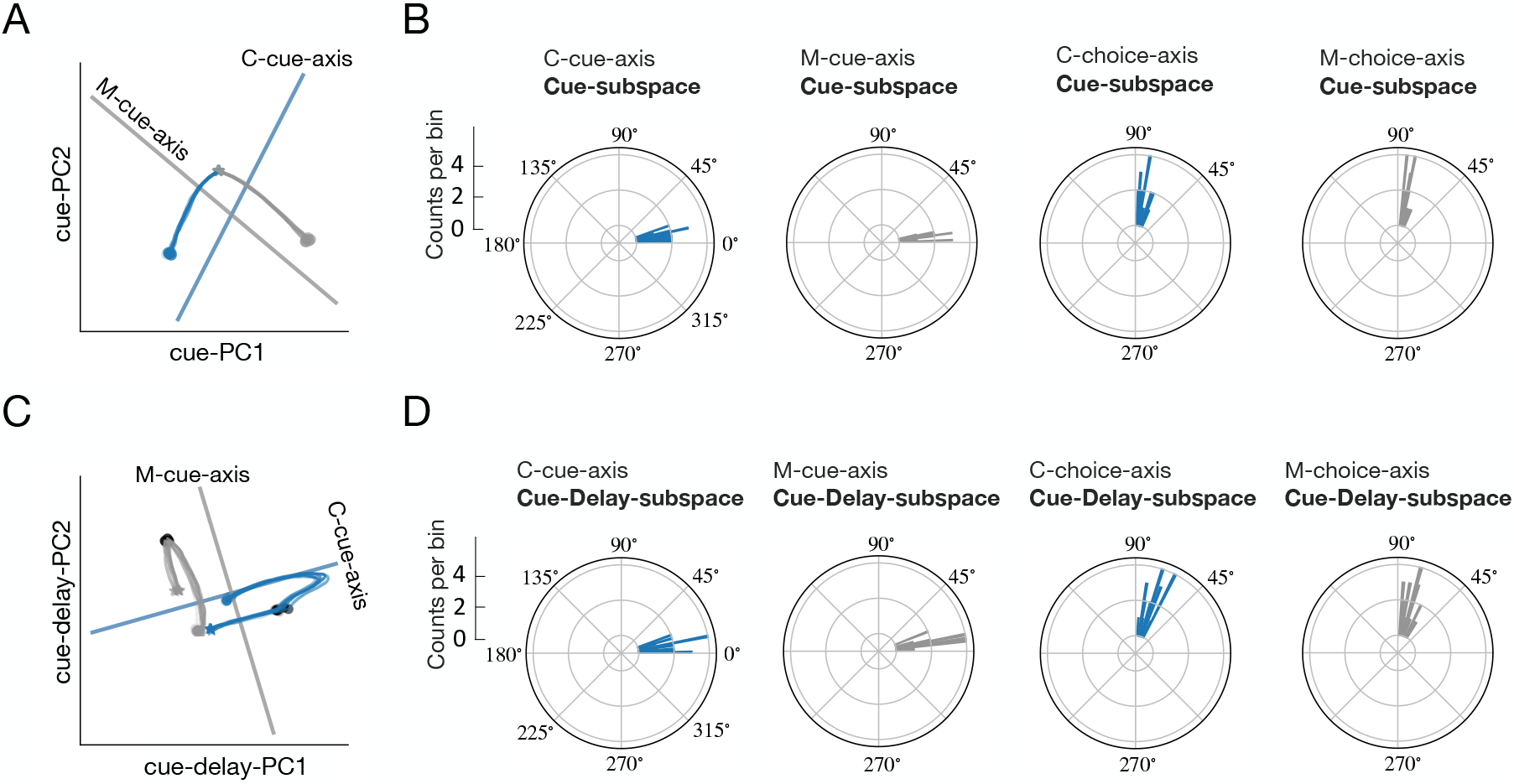
Visualization of Neural Trajectory in Task Subspaces. **(A)** Neural trajectory in the Cue-subspace during the cue stimulus epoch. Blue and grey curves correspond to the color and motion contexts, respectively. Stars and circle indicate the start and end points of the cue stimulus epoch, respectively. Solid lines represent the projections of the `C-cue-axis` (blue) and `M-cue-axis` (grey) in this subspace. **(B)** Polar histograms of the angles between four task-related axes and Cue-subspace during the cue stimulus epoch over 20 training configurations. **(C)** Neural trajectory in the combined Cue-Delay-subspace. Stars denote the neural state at 100 ms after cue stimulus onset, circles denote the beginning points of the cue-delay epoch, and dark circles denote the state at 200 ms before the end of delay. **(D)** Polar histograms of the angles between four task-related axes and Cue-Delay-subspace.

Next, we calculated the angle between Cue-subspace and four predefined axes (*i.e.*, `C-cue-axis, M-cue-axis, C-choice-axis, M-choice-axis`), respectively. Geometrically, if the angle between an axis and a plane is greater than 70°, they are considered nearly orthogonal; if it is less than 20°, they are considered nearly parallel or overlap. We found that the angle between `C-cue-axis` and Cue-subspace, as well as the angle between `M-cue-axis` and Cue-subspace, were both less than 20° (Figure 7*B*). This indicates that the space constructed by the `C-cue-axis` and `M-cue-axis` was overlapping with Cue-subspace. However, the angle between the `C-choice-axis` and Cue-subspace was around 70°-90°, as well as the angle between the `M-choice-axis` and Cue-subspace was both around 70°-90°. Therefore, both the `C-choice-axis` and `M-choice-axis` were nearly orthogonal with Cue-subspace. Based on these observations, we only projected `C-cue-axis` and `M-cue-axis` onto Cue-subspace (Figure 7*A*). We found that in the color context, all trajectories moved along the `C-cue-axis` but were insensitive to the `M-cue-axis`. Similarly, in the motion context, all trajectories moved along the `M-cue-axis` but were insensitive to the `C-cue-axis`. Furthermore, the angle between `M-cue-axis` and `C-cue-axis` mostly centered around 75°-90° (Figure S6*B*), suggesting that these two contextual cues were represented in two almost orthogonal subspaces.

### Cue Information Maintenance

We further explored the question: what is the relationship between neural trajectories across task epochs? The answer to this question can help us understand how the cue information generated in the cue stimulus epoch are preserved during the delay epoch. Specifically, we performed PCA during a 1000 ms time window starting at 100 ms after the cue stimulus presentation and ending at 200 ms before the end of delay epoch. We examined the state trajectory in the two-dimensional space (denoted as Cue-Delay-subspace) spanned by the first two principal components (cue-delay-PC1, cue-delay-PC2), which explained 75% variance (Figure S6*D*). Meanwhile, we projected four predefined axes onto Cue-Delay-subspace (Figure 7*C*), and then calculated the angle between the axes and Cue-Delay-subspace (Figure 7*D*).

Figure 7*C* describes the evolution of the population dynamics in relation to cue stimulus and delay epochs. To depict the transition of population dynamics more clearly, we further divided the 1000-ms time window into two intervals. The first 300-ms interval was within the cue stimulus period, starting from the 100 ms after the cue stimulus onset until the end of cue stimulus. During this interval, the separation between two context-specific trajectories (blue for the color context, grey for the motion context) diverged along their corresponding context cue axes, and reached a maximum at the end of this interval. In the follow-up 700-ms window within the delay epoch, the separation between two context-specific trajectories remained converged along their corresponding context cue axes. Therefore, the separation of context information generated in the cue stimulus epoch was preserved in the delay epoch, even if this separability weakened during the delay epoch. Moreover, for the given context, the projection of trajectories in the `C-cue-axis` at *t* = 100 ms was almost the same as that at *t* = 1000 ms. This correspondence was consistent with the correspondence in the energy of population activity (Figure 3*C*, grey curve).

### Cue-dependent processing of sensory stimulus

Furthermore, we studied how population activity responded appropriately to sensory stimulus according to the current task context. Similarly, we applied PCA to the neural activity during the sensory stimulus epoch. The first three principal components (integ-PCs) explained 81% cross-trial variance (Figure 8*A*), which was caused by the strength and direction of the color evidence, the strength and direction of the motion evidence, and context information (color or motion). As shown by the evolution of the population dynamics in the three-dimensional subspace (Figure 8*B*), each neural trajectory corresponded to a specific task condition, indicating that the activity trajectories in this subspace captured the relationship between the task-related variables.

**Figure 8:**
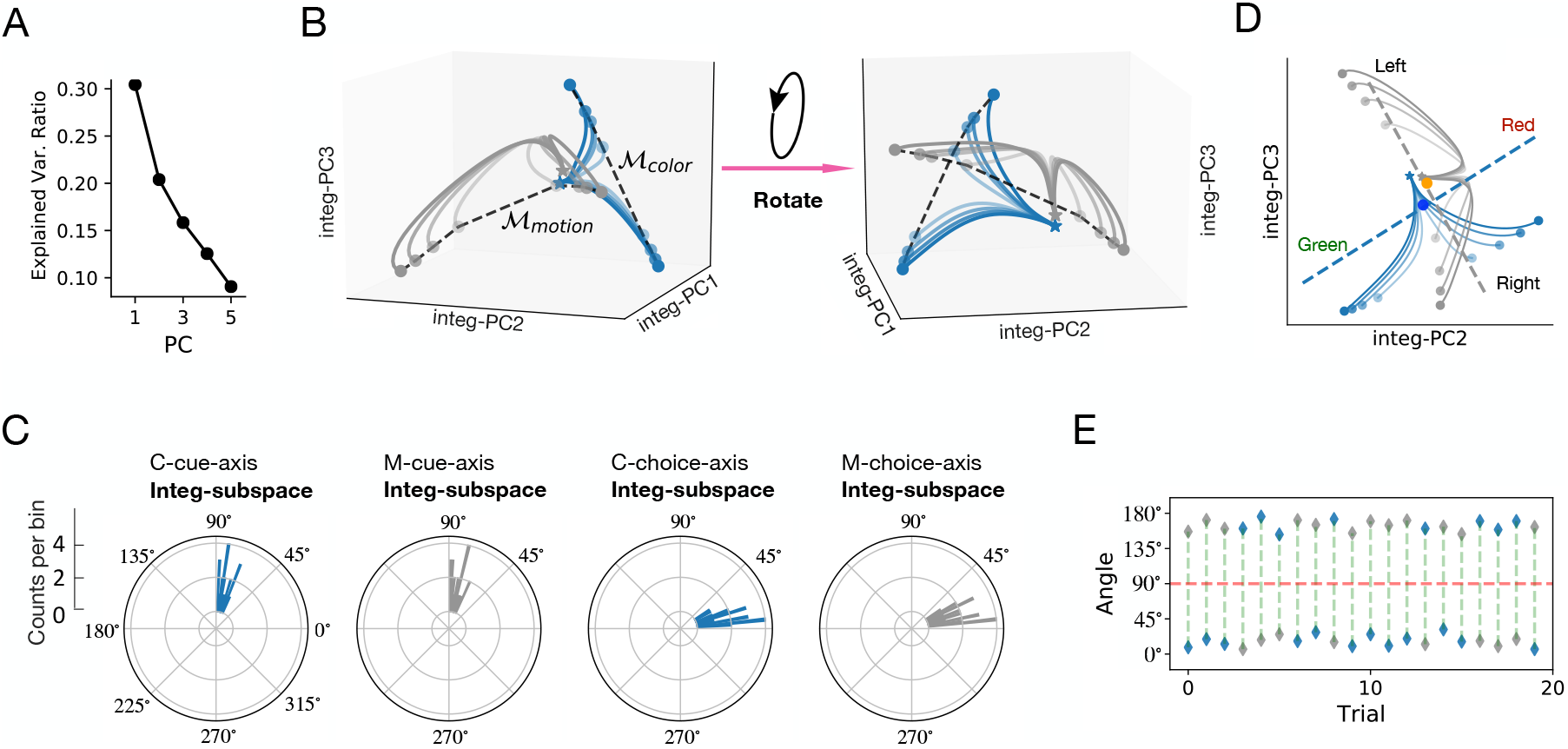
Population Dynamics during the Sensory Stimulus Epoch. **(A)** Ratio of explained variance of the first five PCs in Integ-subspace. **(B)** Neural trajectories in the three-dimensional subspace spanned by integ-PC1, integ-PC2, integ-PC3. **(C)** Polar histograms of the angles between four task-related axes and Integ-subspace. **(D)** Neural trajectories in the two-dimensional subspace spanned by integ-PC2 and integ-PC3. Blue dashed line represents the projection of the `C-cue-axis` and the dark blue solid circle denotes the center point of the projection. Grey dashed line represents the projection of the `M-cue-axis` and the orange solid circle denotes the center point of the projection. **(E)** The angles between integ-PC1 and the `C-cue-axis` (blue) or `M-cue-axis` (grey) in 20 trials.

We further restricted our analysis to the two-dimensional Integ-subspace (spanned by integ-PC2 and integ-PC3). We computed the angle between four task-related axes and Integ-subspace (Figure 8*C*). The angle between the `C-cue-axis` and Integ-subspace, as well as the angle between the `M-cue-axis` and Integ-subspace, both centered around 70°. This means that both the `C-cue-axis` and `M-cue-axis` were orthogonal with Integ-subspace. Moreover, the `C-choice-axis` and `M-choice-axis` were nearly orthogonal (Figure S8*A*). Further, the angle between the `C-choice-axis` and Integ-subspace, as well as the angle between the `M-choice-axis` and Integ-subspace, were both less than 30°. This implies that the subspace spanned by the `C-choice-axis` and `M-choice-axis` was overlapping with Integ-subspace; and the projections of the `C-choice-axis` and `M-choice-axis` onto the Integ-subspace were still orthogonal (Figure 8*D*). Next, we investigated the mechanism of selection and integration through examining the population responses in Integ-subspace in both two-(Figure 8*D*) and three-dimensional subspaces (Figure 8*B*). Several observations are in order.

First, the integration of sensory stimuli corresponded to an evolution of the neural trajectory during the presentation of random dots. All trajectories started from the same starting point in three-dimensional subspace (Figure 8*B*), which corresponded to the initial pattern of population responses during the delay epoch. When the random dots started, the trajectories quickly moved away from their initial state. In Integ-subspace, there was a distinct gradual evolution of the neural trajectory along the `C-choice-axis` and `M-choice-axis` (Figure 8*D*). Specifically, in the color (motion) context trials, the neural trajectory moved along two opposite directions, which corresponded to the two different visual targets, namely, green (left) and red (right).

Second, population responses varied according to different sensory inputs. Specifically, the neural trajectories corresponding to the color context were very different from those corresponding to the motion context, implying that sensory signals were separable at a context-dependent manner. In Figure 8*D*, the blue and grey curves represented neural trajectories in the color and motion contexts, respectively. In the color context, the patterns of population responses also varied with respect to different strengths and direction of the color stimulus. Therefore, neural trajectories captured multi-dimensional task-related variables, such as the context information, strength, and direction of sensory stimulus. Moreover, instead of following a straight line along the `C-choice-axis` or `M-choice-axis`, the neural trajectory formed an arc within the corresponding choice axis. The distance between the projection point of each arc onto the `C-choice-axis` and the color-center point (dark blue circle) reflected the strength of the corresponding color evidence. While the position (two sides of the color-center point) of the projection point of each arc onto the color axis reflected the direction of the target (towards green or red). Once the random points disappeared, the network stopped to integrate the sensory evidence, yet the integrated evidence continued to be preserved along the `C-choice-axis` (Figure 9*A,B*). Similar discussions also held in the motion context.

**Figure 9:**
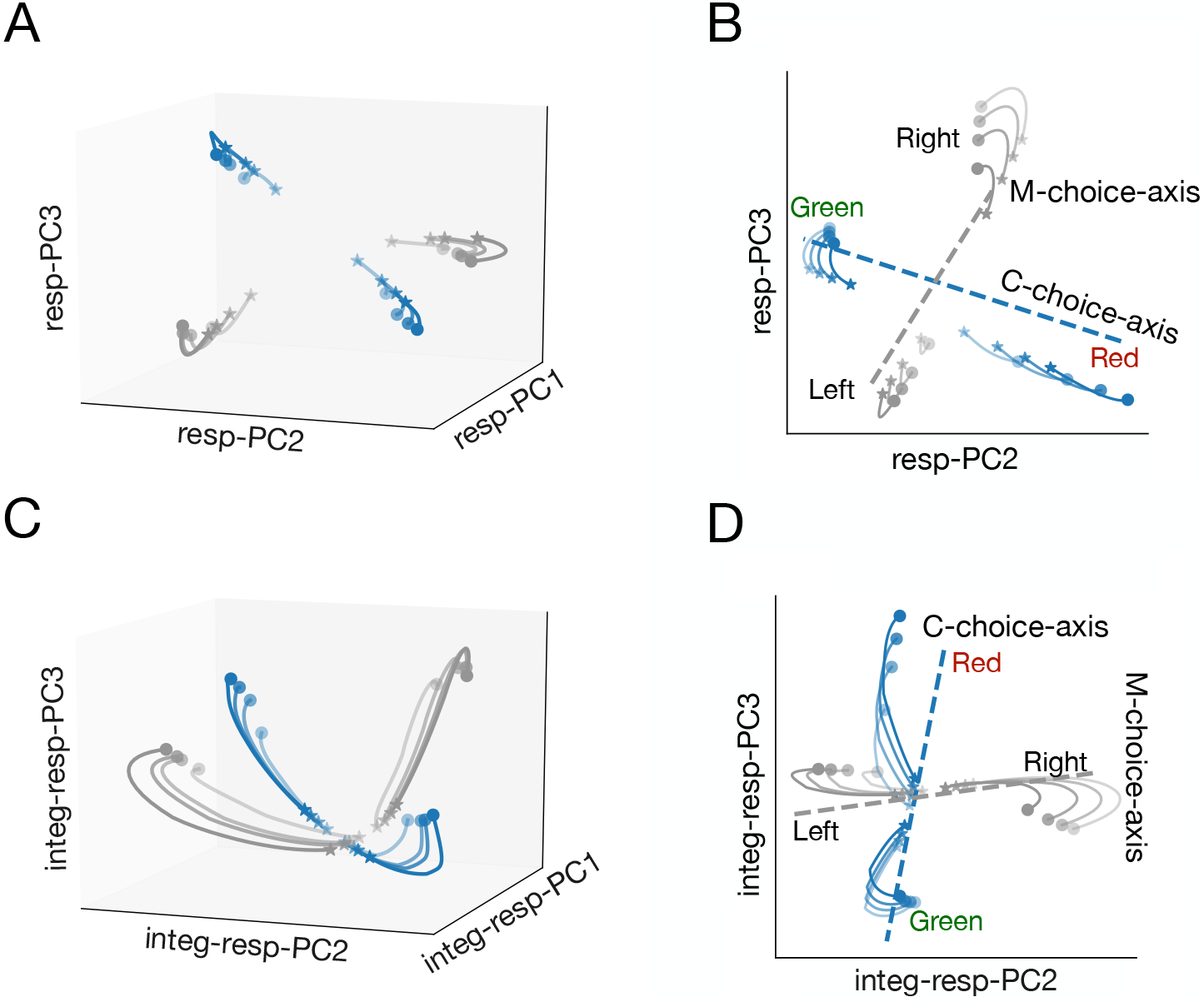
Visualization of Neural Trajectory in the Response Subspace. **(A)** Neural trajectories in the three-dimensional subspace spanned by resp-PC1, resp-PC2, resp-PC3 during the response epoch. **(B)** Same as panel A, except in two-dimensional subspace. **(C)** Neural trajectories in the three-dimensional subspace spanned by integ-resp-PC2, integ-resp-PC2, integ-resp-PC3. Each trajectory started from 500 ms after the beginning of sensory stimulus until the action response. **(D)** Same as panel C, except in two-dimensional subspace.

Third, population responses in the color and motion contexts occupied two different parts of subspace, which corresponded to two orthogonal-but-interesected subspaces. According to the above discussion, the neural trajectory in the color context was guided by the `C-cue-axis` and `C-choice-axis`. Therefore, the population activity during the color context occupied a subspace (*i.e.*, Color-subspace spanned by the `C-cue-axis` and `C-choice-axis`). In a similar fashion, population activity in the motion context occupied the Motion-subspace spanned by the `M-cue-axis` and `M-choice-axis`. We found that the projection of all neural trajectories onto an irrelevant choice axis was almost the same at each moment (Figure 8*D*). This indicates that the Color-subspace and Motion-subspace were mutually orthogonal: the sensory information in the color context couldn’t be captured by the `M-choice-axis`, and the sensory information in the motor context couldn’t be captured by the `C-choice-axis`. Moreover, we calculated the angle between the `C-cue-axis` and the integ-PC1, as well as the angle between the `M-cue-axis` and the integ-PC1. Noting that the range of this angle was 0° ∼ 180°, which helped us detect whether the projection directions of the two cue axes on integ-PC1 were opposite. We found at each trial, one angle was greater than 90°, and the other was less than 90° (Figure 8*E*). That is, the projection directions of the `C-cue-axis` and `M-cue-axis` on integ-PC1 were always opposite, suggesting that Color-subspace and Motion-subspace were orthogonal but intersected.

### Task Response

We applied PCA to the population activity during the response epoch, and found that the first three principal components (resp-PCs) explained 96% data variance (Figure S10*A*). The three-dimensional trajectories under different trial conditions occupied different positions in the Resp-subspace (Figure 9*A*). Similarly, we calculated the angle between the four task-related axes and the Resp-subspace (spanned by resp-PC2, resp-PC3). The result showed that: (i) The angle between `C-cue-axis` and Resp-subspace, as well as the angle between `C-cue-axis` and Resp-subspace, were both around 70°-90°. (ii) The angle between the `C-choice-axis` and Resp-subspace, as well as the angle between the `M-choice-axis` and Resp-subspace, both centered around 30° (Figure S10*B*). Therefore, the near orthogo-nality between `C-choice-axis` and `M-choice-axis` could be maintained in Resp-subspace (Figure 9*B*). (iii) The projection of the neural trajectory on the corresponding choice axis was almost the same at each moment, suggesting that there was no sensory evidence integration during the response epoch (Figure 9*B*).

To examine how the neural trajectory evolved from the sensory stimulus epoch to the response epoch, we reapplied PCA to the population activity during a combined-epoch 600-ms period: starting at 500 ms after the presentation of the sensory stimulus and ending at the moment of decision. The first three PCs (integ-resp-PCs) explains 92% cross-trial variance (Figure S11*B*). The neural trajectory in the two-dimensional subspace (denoted as Integ-Resp-subspace, Figure 9*D*) had similar characteristics to that during the sensory stimulus period: First, the angle between the `C-choice-axis` and Integ-Resp-subspace, as well as the angle between the `M-choice-axis` and Integ-Resp-subspace, both centered around 30° (Figure S11*A*). Second, the projection directions of the `C-cue-axis` and `M-cue-axis` on integ-resp-PC1 were always opposite (Figure S11*C*), thus the integ-resp-PC1 fully captured the context cue information. Third, the projection of all neural trajectories on an irrelevant choice axis was the same at each moment (Figure 9*D*). Therefore, the population activity during this extended period also occupied two orthogonal-but-intersected subspaces.

Finally, we investigated the impact of weight perturbation on low-dimensional neural trajectories. Specifically, we perturbed the recurrent connection weights and examined the neural activity trajectory in the corresponding three-dimensional subspace. The range of local and global perturbation was restricted, and we examined the neural trajectories during the delay epoch and sensory stimulus epoch, respectively. Under a small weight perturbation, the relationship between task-related variables, such as the strength and direction of the random dots, remained relatively robust (Figure S7 and Figure S9). When the level of perturbation increased, the task-related information became lost gradually.

### A Geometric Interpretation of Context-Dependent Integration

Based upon our geometric framework and above subspace analyses, we propose a geometric interpretation for context-dependent computation during WM and decision making. To further illustrate the geometric notion, we defined dynamic attractors (such as fixed points) in the high-dimensional RNN dynamics (Sussillo and Barak, 2013).

Specifically, we used numerical optimization to minimize 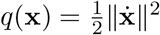 to identify the fixed points and slow points. Each dot in Figure 10*A* represents a slow point with *q*(**x**) < 0.01, and each cross represents a fixed point with *q*(**x**) < 0.0001. These fixed points were further arranged along a line for a specific context (blue crosses: color context; black crosses: motion context), forming a line attractor (Figures 10*A* and 10*B*).

**Figure 10:**
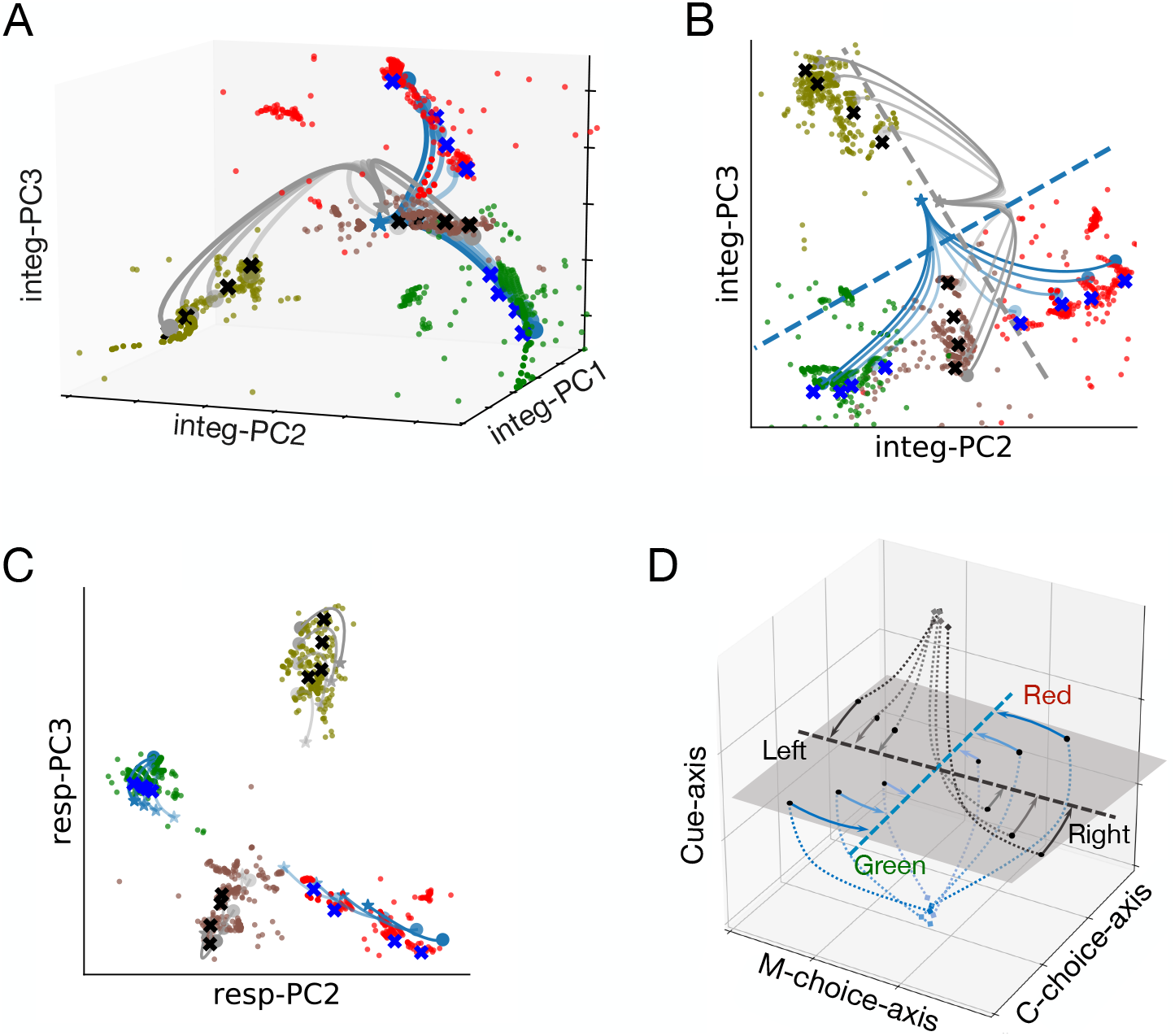
Visualization of Dynamic Attractors in Neural Subspace. (A) Neural trajectories in three-dimensional Integ-subspace for different sensory stimuli eventually converged to different attractors. Blue and black crosses correspond to the color and motion contexts, respectively. Dots denote slow points, and different colors correspond to different trial stimuli (red dots as red stimuli, green dots as green stimuli, olive dots as leftward direction stimuli, and brown dots as rightward direction stimuli). **(B)** The projection of slow points and fixed points onto Integ-subspace. **(C)** The projection of slow points and fixed points onto the Resp-subspace. **(D)** A schematic of neural trajectory through a three-dimensional subspace during the transition from evidence of sensory integration to choice making. Three axes include two choice axes and one context cue axis. Dotted lines represent neural trajectories during the sensory stimulus epoch, and solid lines reflect neural trajectories during a 75-ms window following the saccade onset.

We have showed the end of neural trajectories during the sensory integration epoch were aligned in manifold 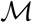 (Figure 8*B*); and the projection of manifold 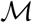 in the Integ-subspace was parallel to the corresponding choice axis (Figure 8*D*), suggesting that the color and motion information could be captured by manifold 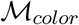 and manifold 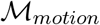, respectively. Interestingly, the line attractors were aligned near the manifold, suggesting that the integration of sensory evidence could be explained by the arrangement of fixed-points: During the sensory integration epoch, the population dynamics drove context-specific trajectories to back to their attractors associated with the correct choice. That is, the population activity was attracted toward the corresponding manifold 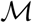 of slow dynamics at the end of the sensory stimuli (Figure 8*B*). By projecting these slow points and fixed points onto Resp-subspace, the population trajectories evolved around the corresponding fixed-points (Figure 10*C*). Additionally, the sensory information integrated during the sensory stimuli epoch was maintained during the response epoch.

Moreover, our analysis within the proposed geometric framework has shown that Color-subspace and Motion-subspace were orthogonal-but-intersected. Therefore, the color contextual cue triggered a dynamical process so that the relevant color evidence was integrated, while ignoring the irrelevant motion evidence from the same trials because of their mutual orthogonality in the subspace. The opposite pattern was also evident in the motion context.

## DISCUSSION

Context-dependent computation is an important hallmark for achieving cognitive flexibility. However, the computational principles underlying context-dependent WM or decision making remains incompletely understood. We trained an RNN to perform a delayed context-dependent decision-making task, and proposed a geometric framework that helps uncover population dynamics of the trained RNN. Importantly, the trained RNN produced some emergent neurophysiological features at both single unit and population levels. The PCA and weight perturbation analysis further revealed neural representations of context-specific dynamic population coding and information integration. In low-dimensional neural subspaces, the RNN encoded the context information through the separation of neural trajectories and maintained the context information during the delay epoch. Finally, sensory integration during the decision-making period can be viewed by an evolving neural trajectory that occupies in two orthogonal-but-intersected subspaces.

Artificial RNNs are generally considered as black-box models, and the underlying dynamical mechanisms are poorly understood. To unravel the black box, Sussillo and Barak (2013) proposed a dynamical systems framework to uncover the computational mechanisms of RNNs. In a similar manner, we used reverse-engineering and subspace analyses to uncover the mechanisms of population coding in the trained RNN. At the single unit level, many units exhibited mixed-selectivity to different task-related variables. At the population level, sequential activation (‘neural sequences’) and rotational dynamics emerged from the trained RNN during the delay period. The neural sequential activation has been found in the mouse prefrontal or parietal cortices during several behavioral tasks (Harvey et al., 2012; Schmitt et al., 2017). Our results of neural sequences and SI were also consistent with the recent report based on RNN modeling (Orhan and Ma, 2019; Bi and Zhou, 2020). Dimensionality reduction-based subspace analyses have been widely used to uncover temporal dynamics of RNNs (Kobak et al., 2016; Kao, 2019), geometric notions can further reveal the orthgonality of task variables in the neural subspace (Bi and Zhou, 2020).

Furthermore, our analyses demonstrated that the context cue triggered the separation of neural trajectories in a low-dimensional subspace, and the dynamic system was in a high-activity state. During the delay epoch, the population activity decayed to a stable low-activity state while maintaining the context separation. During the random dots presentation, we found two main features of population responses: (i) The population responses exhibited different patterns to varying sensory inputs. (ii) The population response in the color and motion contexts occupied two orthogonal-but-intersected subspaces. These features of population responses can be summarized schematically in Figure 10*D*, which provide fundamental constraints on the mechanisms of context-dependent computation in flexible cognitive tasks.

### RNNs for Understanding Computational Mechanisms of Brain Functions

Due to its powerful computational capabilities, RNNs exhibit complex dynamics similar to experimental findings (Wolfgang et al., 2002; Sussillo and Abbott, 2009; Buonomano et al., 2009; Mante et al., 2013). In an analogue to animal behavioral training, RNNs can perform a wide range of cognitive tasks with supervised learning and labeled examples, including WM (Barak et al., 2013b, 2010; Rajan et al., 2016), motor control (Laje and Buonomano, 2013; Hennequin et al., 2014), and decision-making (Song et al., 2016; Sussillo et al., 2015; Thomas, 2017; Yang et al., 2019). The activity and network connectivity of the trained RNN can be accessed with specific perturbation strategies to help reveal underlying computational mechanisms. For example, Chaisangmongkon et al., (2017) discovered the features of structural and functional connectivity that support robust transient activities by training RNN to solve a match-to-category task, and the trained network could capture various phenomena from neurophysiological experiments. Orhan and Ma (2019) identified the circuit-related and task-related factor that generate the sequential or persistent activity by training the RNN to perform various WM tasks. Goudar and Buonomano (2018) demonstrated that time-varying sensory and motor patterns can be stored as neural trajectories within the RNN, helping us understand the time-warping codes in the brain. Additionally, some RNN models with more basic biological features, such as Dale’s principle, have been developed as a valuable platform for generating or testing new hypotheses (Song et al., 2016). An excitatory-inhibitory spiking RNN that incorporates more biological constraints, such as spike frequency adaptation, has been used to learn a context-dependent working memory task (Xue et al., 2021).

### Relation to Other Working Memory Theories

In context-dependent computational tasks, to decide which type of subsequent sensory inputs to be integrated, the information about the context rule needs to be preserved as working memory across different task epochs. In the past, various theories and computational models for WM have been developed. The classic models suggest that the PFC support WM via the stable, persistent activity of stimulus-specific neurons (Wang, 2001; Miller and Cohen, 2001; Miller, 2000), which bridges the gap between the memory representation of the context cue and sensory stimulus epochs. However, some experimental observations shows that neural population coding in WM is dynamic over the course of a trial (Barak et al., 2010; Stokes et al., 2013). Classical and new theoretical models in WM have been reviewed in light of recent experimental findings (Lundqvist et al., 2016; Miller et al., 2018). Similar to other modeling effort (Orhan and Ma, 2019; Xue et al., 2021), we found that the single-unit response in WM showed strong temporal dynamics rather than persistent activity. Moreover, unlike previous studies that suggested a dissociation between the stimulus-driven response and subsequent delay activity in the PFC (Barak et al., 2010; Meyers et al., 2008), our results showed the opposite phenomenon. In fact, the context information encoded by population activity during the cue stimulus epoch was maintained by low energy activity (Figure 3*C*) and sequential representation (Figure 4*F*) during the delay epoch.

Population coding is highly dynamic during both the cue stimulus and the delay epoch (Figure 3*C*). One possible explanation for the dynamic activity in the early part of the delay epoch is that it reflects the transformation from a transient context cue input into a stable WM representation, and this transformation is first contained in a high-energy dynamic trajectory in the state space, and then the system returns to a low-energy state. Moreover, Mongillo et al. (2008) have proposed a computational model of WM, in which the memories can be maintained as a pattern of synaptic weights. Specifically, neural activity changes synaptic efficacy within their computational model, leaving a synaptic memory trace via short-term synaptic plasticity. This suggests that the previous cue stimuli may be recovered from the network architecture, allowing for an energy-saving mode of short-term memory, rather than relying on the maintenance of high-energy persistent activity. Based on this synaptic theory, Stokes (2015) have predicted that WM representations should be stationary and have low energy. Our analysis results are also consistent with this prediction.

### Computational Mechanism Revealed by Analyses of Population Activity

In many brain regions, information is encoded by the activity patterns of population. Thinking of the neural population activity as states in a dynamical system is increasingly prevalent in neuroscience (Sauerbrei et al., 2020; Shreya et al., 2020; Churchland et al., 2012). The population activity over time corresponding to trajectories in the neural space, and visualization of state trajectories allows us to detect the population dynamics. Dimensionality reduction techniques have been developed to find dimensions relevant to dynamical structure, which identify a low-dimensional subspace for population responses (Churchland et al., 2012; Kobak et al., 2016). Finally, although we have focused our RNN modeling on a cognitive task, our geometric framework and subspace analysis can be applied to investigate other brain areas or brain functions, such as the premotor cortex or primary motor cortex in various motor tasks (Elsayed et al., 2016; Kao, 2019; Sussillo et al., 2015).

## STAR Methods

Detailed methods include the following:

- KEY RESOURCE TABLE
- CONTACT FOR REAGENT AND RESOURCE SHARING
- METHOD DETAILS

– Network Structure
– Task Description
– RNN Training
– Definition of Task-Related Variables
- QUANTIFICATION AND STATISTICAL ANALYSIS

– Computation of Sequentiality Index (SI)
– Finding Rotation Dynamics via jPCA
– Finding Fixed Points and Line Attractor
- DATA AND SOFTWARE AVAILABILITY

## KEY RESOURCE TABLE

| REAGENT or RESOURCE | SOURCE | IDENTIFIER |
| --- | --- | --- |
| Software and Algorithm |  |  |
| Python | Open source | www.python.org |
| Sequentiality index | Orhan and Ma (2019) | https://github.com/eminorhan/recurrent-memory |
| Fixed point analysis | Sussillo and Barak (2013) | https://github.com/elipollock/EMPJ |
| Network structure | Yang et al. (2019) | https://github.com/gyyang/multitask |
| Dimensionality reduction and sequence visualization | Bi and Zhou (2020) | https://github.com/zedongbi/IntervalTiming |
| Deposited code | This study |  |

## CONTACT FOR REAGENT AND RESOURCE SHARING

Further information and requests for data should be directed to and will be fulfilled by the Lead Contact, Zhe S. Chen.

## METHOD DETAILS

### Network Structure

We constructed an RNN network of *N* = 256 fully interconnected neurons described by a standard firing-rate model. The continuous-time formulation dynamics of the RNN are govern by following equations

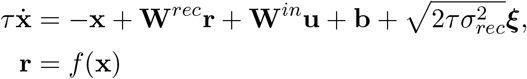

where **x**, **r**, and **u** represent the synaptic current, firing rate and network input, respectively; *τ* = 20 ms is time constant, which mimics the synaptic dynamic on the basis of NMDA receptors; **b** is the background input; ****ξ**** are independent Gaussian white noise scaled by *σ*_*rec*_ = 0.05, which represent the intrinsic noise. The firing rate **r** is related to the corresponding current **x** by a softplus transfer function *f* (*x*) = log(1 + exp(*x*)), which maps every input currents to a positive firing rate. The output of the network is a linear mapping between the firing rate and a readout synaptic weight:

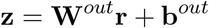

where **W**^*in*^, **W**^*rec*^, **W**^*out*^ are the input weight, recurrent weight and output weight, respectively. We used Euler’s method to discretize the continuous-time equation and derive the discrete-time version

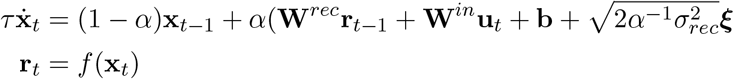

where 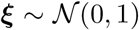 was drawn from a standard normal distribution; *α* = Δ*t/τ*, and Δ*t* is time step. In our study, we set Δ*t* = 20 ms. Similar to the previous computational models (Wang et al., 2018; Orhan and Ma, 2019), we used *α* = 1. Furthermore, we assumed that the RNN received two types of noisy input: rule-specific input **u**_*rule*_ and stimulus-specific input **u**_*stim*_:

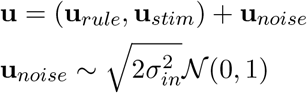

where *σ*_*in*_ denotes the standard deviation of input noise. We set *σ*_*rec*_ = 0.05, *σ*_*in*_ = 0.01 for training the RNN.

### Task Description

The task is schematized in Figure 1*A*, which is inspired from context-dependent task performed by macaque monkeys (Mante et al., 2013). The network received pulses from three input channels. The first channel consisted of the context cue stimulus, which contained two units encoding different context information. The other two channels consisted of two different sensory stimuli modalities. One channel represented the color sensory stimulus, which contained two units encoding the color stimulus strength *γ*_*color,*1_, *γ*_*color,*2_. The other channel represented the motion sensory stimulus, which encoded the motion stimulus strength *γ*_*motion,*1_, *γ*_*motion,*2_. The stimulus strengths were determined by the coherence for the color modality and motion modality (*c*_*color*_, *c*_*motion*_), and set as follows:

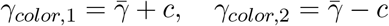

A similar equation held for the motion modality. 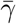 denotes the average strength of the two color stimuli, which was drawn from a uniform distribution 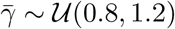 (where 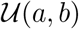 represents a uniform distribution between *a* and *b*). Coherence *c* measured the strength difference of these two stimuli, which was uniformly distributed as

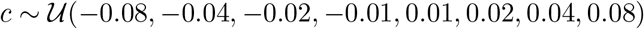

The task consisted of distinct epochs. For each trial, a fixation epoch was present before the stimulus presentation. It was followed by the context cue stimulus epoch that lasted *T*_*stim*1_ = 400 ms. After a delay epoch (with duration of *T*_*delay*_ = 800 ms), random dots stimulus was presented in the second stimulus epoch with a duration of *T*_*stim*2_ = 800 ms. Finally, the network responded in the Go epoch with an interval of *T*_*resp*_.

### RNN Training

If the relevant evidence points towards choice 1, the output channel 1 (composed by two output units) was activated, otherwise the output channel 2 (composed by two output units) was activated. The cost function *L* is the mean squared error (MSE) between the network output (**z**) and target outputs 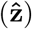:

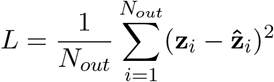

We optimized the weights {**W**^*in*^, **W**^*rec*^, **W**^*out*^} using the well-established Adam algorithm (Kingma and Ba, 2015), with default configuration of hyperparameters. The learning rate is 0.0005, and the exponential decay rate for the first and second moment estimates are 0.9 and 0.999, respectively. The off-diagonal connections of recurrent weight matrix **W**^*rec*^ were initialized using a normalized random Gaussian matrix, and diagonal connections were initialized to 1. The initial input connection weights were uniformly drawn from −0.5 to 0.5. The output connection weights **W**^*out*^ were initialized as independent Gaussian random variable with mean 0, and standard deviations 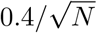.

### Definition of Task-Related Axes

We first grouped neural population activity into the matrix **X**∈ ℝ^*N* ×*CT*^, where *N* = 256 denotes the number of RNN units, *C* = 1 × 8 denotes the number of the conditions (8 different sensory stimulus conditions in the given context), and *T* denotes the time step.

Spike activity was binned by a 20-ms window. Different task epochs correspond to different **X**-matrices.

`C-cue-axis` and `M-cue-axis`. During the cue stimulus epoch, for the given color context, we obtained the matrix **X**_*cue,color*_ ∈ ℝ^*N* ×*CT*^, where *N* = 256, *T* = 400*/*20. We further performed PCA on the matrix **X**_*cue,color*_. The first PC explained 91% of data variance (Figure S6*C*, green bar), so we defined it as the `C-cue-axis`. Similarly, for the given motion context, we performed PCA on the matrix **X**_*cue,motion*_ ∈ *R*^N×*CT*^ and the ratio of explained variance of the first five PCs is also shown in Figure S6*C* (grey bar). The first PC explained 92% of data variance. Therefore, we defined the first PC dimension as the `M-cue-axis`(Table 1).

`C-choice-axis` and `M-choice-axis`. The population activity matrices in the given context were **X**_*integ,color*_ ∈ ℝ^*N* ×*CT*^ and **X**_*integ,motion*_ ∈ ℝ^*N* ×*CT*^, where *T* = 800*/*20. We performed PCA on **X**_*integ,color*_ ∈ ℝ^*N* ×*CT*^ and **X**_*integ,motion*_ ∈ ℝ^*N* ×*CT*^, respectively. The ratio of explained variance of the first five PCs in the specific context is shown in Figure S8*B*. In a similar fashion, we defined `C-choice-axis` and `M-choice-axis` for the color and motion contexts, respectively.

## QUANTIFICATION AND STATISTICAL ANALYSIS

### Computation of Sequentiality Index (SI)

We computed the SI to quantify the sequential activation of the population response of excitatory neurons during the delay epoch. The SI is defined as the sum of the entropy of the peak response time distribution of the recurrent neurons and the mean log ridge-to-background ratio of the neurons, where the ridge-to-background ratio for a given neuron is defined as the mean activity of the neuron inside a small window around its peak response time divided by its mean activity outside this window (Orhan and Ma, 2019).

### Finding Rotation Dynamics via jPCA

The jPCA method has been used to reveal rotational dynamics of neuronal population responses (Churchland et al., 2012). We assumed that the data were modeled as a linear time-invariant continuous dynamical system of the form: 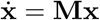, where the linear transformation matrix **M** was constrained to be skew-symmetric (*i.e.*, **M**^T^ = −**M**). The jPCA algorithm projects high-dimensional data **x**(*t*) onto the eigenvectors of the **M** matrix, and these eigenvectors arise in complex conjugate pairs. Given a pair of eigenvectors 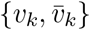, the *k*-th jPCA projection plane axes are defined as 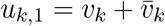 and 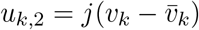 (where 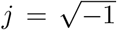). The solution to the above continuous-time differential equation is given by **x**(*t*) = *e*^**M***t*^**x**(0), where the family {*e*^**M***t*^} is often referred to as the semi-group generated by the linear operator **M**. Since **M** is skew-symmetric, *e*^**M**^is orthogonal; therefore, it can describe the rotation of the initial condition **x**(0) over time.

Applying eigenvalue decomposition to the real skew-symmetric matrix **M**, so that **M** = **UΛU**^−1^, where **Λ** is a diagonal matrix whose entries 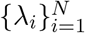 are a set of (zero or purely imaginary) eigenvalues. Upon time discretization (assuming *dt* = 1), we obtained the discrete analog of dynamic equation **x**(*t* + 1) = (**I**+ **M**)**x**(*t*). Alternatively, we directly solved a discrete dynamical system of the vector autoregressive (VAR) process form **x**(*t*+1) = **Qx**(*t*) over the space of orthogonal **Q**matrices. Mathematically, we have previously shown that this is equivalent to solving the following constrained optimization problem (Nemati et al., 2014):

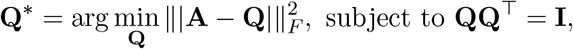

where || · ||_*F*_ denotes the matrix Frobenius norm, and 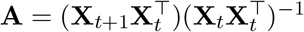 represents the least square solution to the unconstrained problem **x**(*t* + 1) = **Ax**(*t*). The solution to the above constrained optimization is given by the orthogonal matrix factor of Polar Decomposition of matrix **A**, namely **A**= **QP**(Higham, 1986).

### Finding Fixed Points and Line Attractor

We focused on finding the fixed points and slow points of dynamical system to explore the mechanism by which the RNN performed the context-dependent task. The computational task was to find some points that satisfy 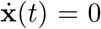 for all *t*, that is, we needed to solve a first-order differential equation

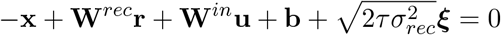

for a constant input **u** and with a transfer function **r** = *f* (**x**). However, the nonlinear function *f* (**x**) makes it difficult to find an analytical solution to differential equation. Therefore, we identified the fixed points and slow points through numerical optimization. According to the algorithm described in Sussillo and Barak (2013), we solved the optimization problem as follows

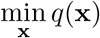

where 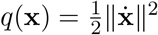. To identify the local minimum, we defined the RNN state **x** as a slow point if *q*(**x**) < 0.01, and **x** is a fixed point if *q*(**x**) < 0.0001. These fixed points and slow points were calculated during the sensory stimulus epoch. The initial conditions for optimization were points in the neighborhood of **x**(*t*), which was the start state of RNN system trajectories. For each fixed point, we repeated the optimization procedure 300 times, and the initial conditions at each time were sampled form the neighborhood of **x**(*t*). Based on these 300 candidate fixed points, we choose stable fixed points as attractors, characterized by slow dynamics. We determined a fixed point being stable if neural states empirically converged to attractors. Here, we only considered stable fixed points (represented by cross symbols in Figure 10). For a given context, there were 8 different trial conditions, and the population activity on each trial condition was characterized by a state trajectory (Figure 8*C*). In total, we identified 16 attractors and each state trajectory eventually converged to its fixed-point attractor (Figure 10*A*). The initial conditions for slow point optimization were sampled from a normalized random Gaussian matrix. We repeated the optimization procedure 56 times for each trial condition and obtained 56 × 16 = 896 slow points.

## DATA AND SOFTWARE AVAILABILITY

Data and code are available upon request to the Lead Contact.

## ACKNOWLEDGMENTS

This work was supported by the a grant (No. 11172103) from the National Natural Science Foundation of China (S.L.).

## AUTHOR CONTRIBUTION

X.Z., S.L. and Z.S.C. designed the experiment. X.Z. performed all experiments and analyses. X.Z. and Z.S.C. wrote the manuscript.

## Supplemental Information

**Figure S1:**
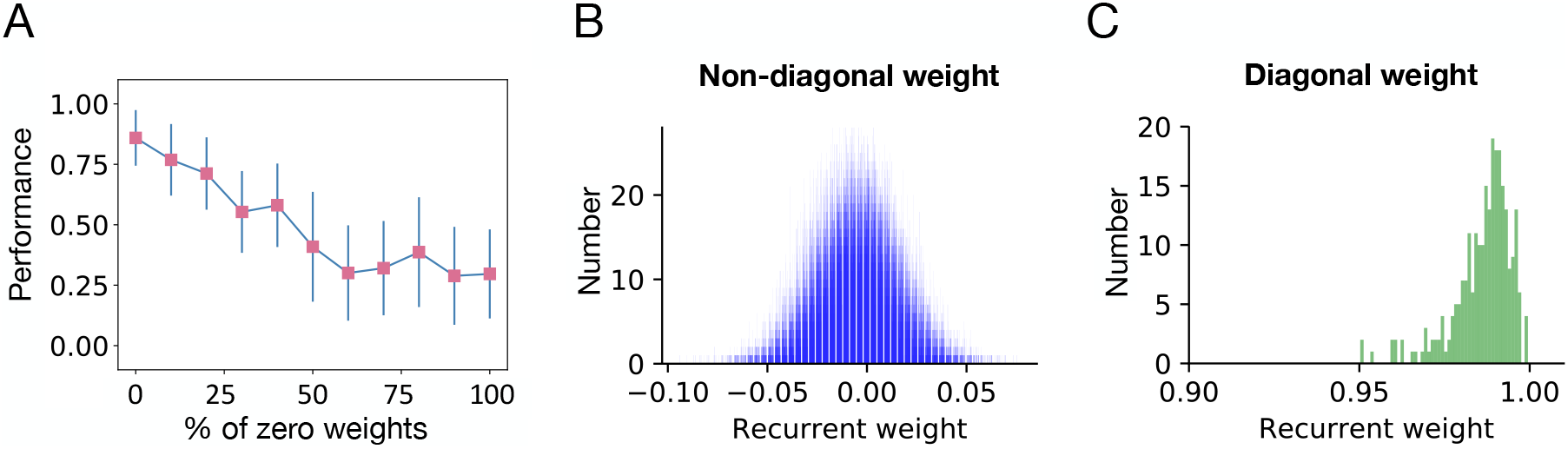
Distributions of recurrent connection weights of trained RNN and the impact of the sparsity on the RNN performance.**(A)** The accuracy of RNN performance with respect to the percentage of zero weight entries. **(B)** Histogram of non-diagonal recurrent weight connections of the trained RNN. **(C)** Histogram of diagonal self-connection weights of the trained RNN.

**Figure S2:**
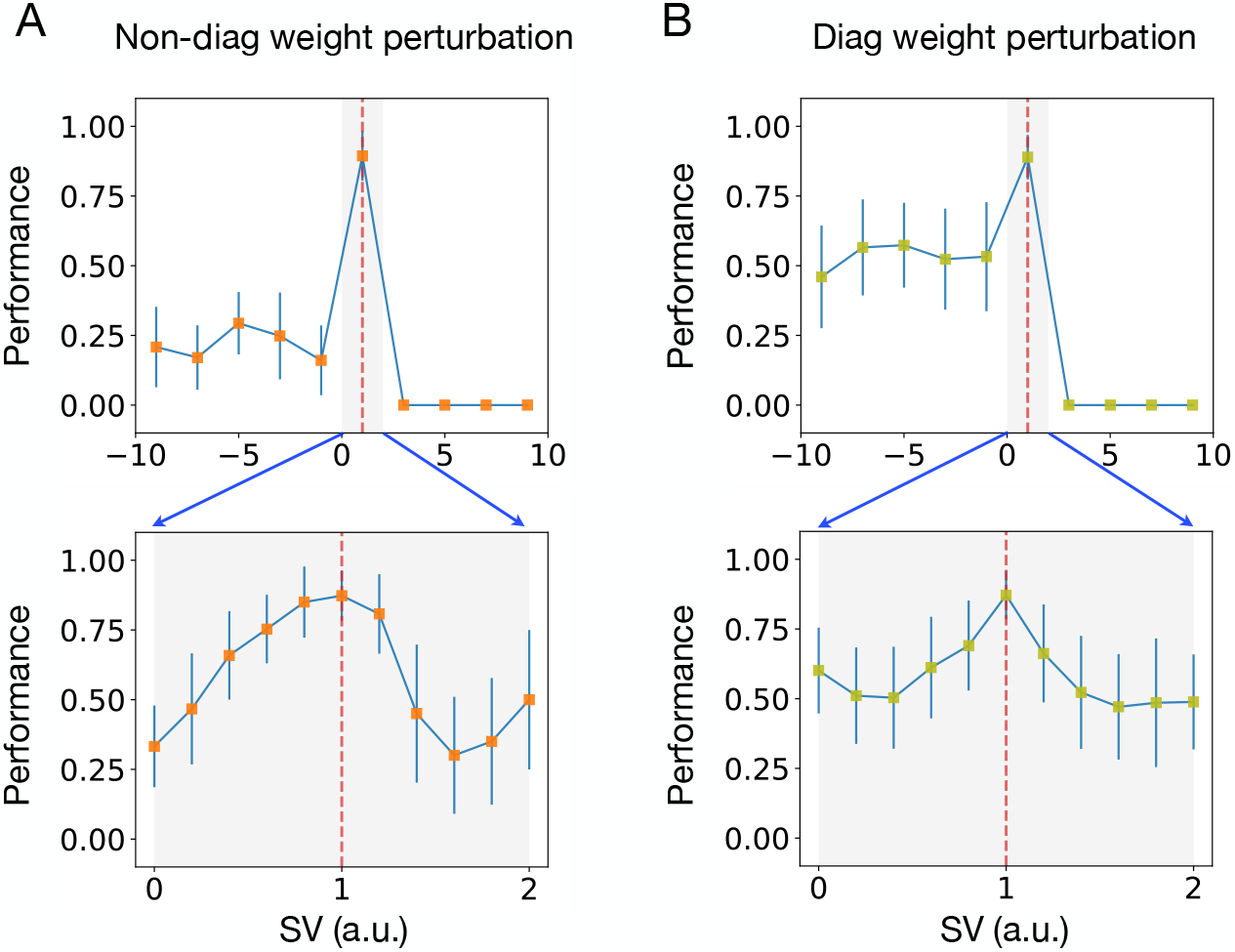
Performance of the trained RNN with respect to recurrent weight perturbation.Perturbing the trained RNN by scaling the recurrent connection weights with different scale values (SVs). **(A)** The impact of non-diagonal weight perturbation on the performance. The red vertical dashed line corresponds to the non-perturbed RNN model (*SV* = 1). The error bars indicate SEM over 20 trained models. The zoom-in shade area is shown in the bottom panel. The trained RNN preserved a robust performance between *SV* = 0 and *SV* = 1.2. **(B)** Similar to panel A, but for diagonal weight perturbation. The trained RNN preserved a robust performance between *SV* = 0.9 and *SV* = 1.1.

**Figure S3:**
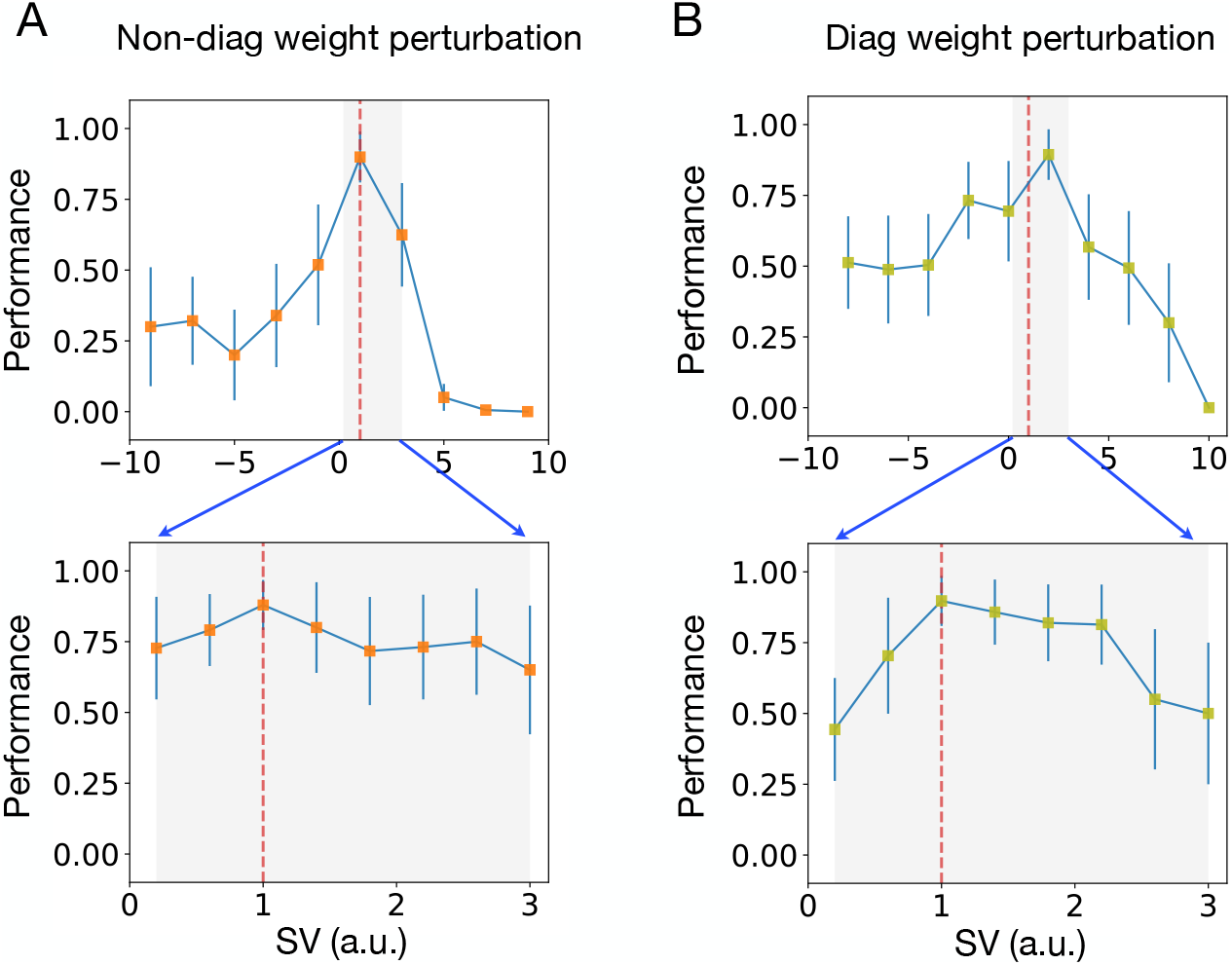
Perturbing the trained RNN by scaling weights locally in time during the delay epoch. **(A)** The top panel plots the impact of non-diagonal connection weight perturbation on the performance. The network showed a robust performance between *SV* = 0.5 and *SV* = 1.5. **(B)** The impact of diagonal recurrent weight perturbation on the performance. The network showed a robust performance between *SV* = 1.01 and *SV* = 2.

**Figure S4:**
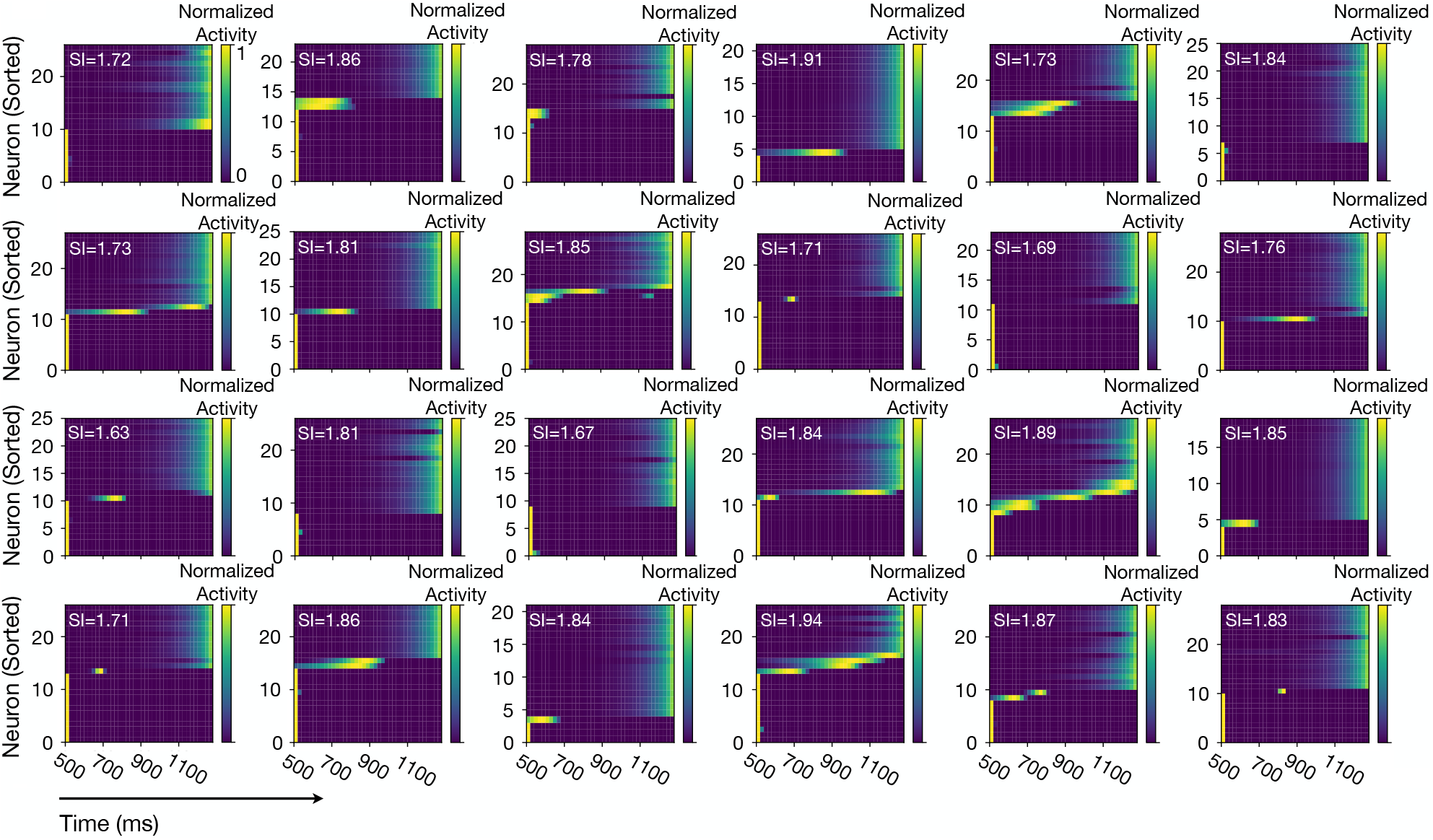
Selected 24 heat maps of sorted population response from 30 randomly selected units. In each plot, 30 units were normalized by their maximal responses and sorted by the peak time.

**Figure S5:**
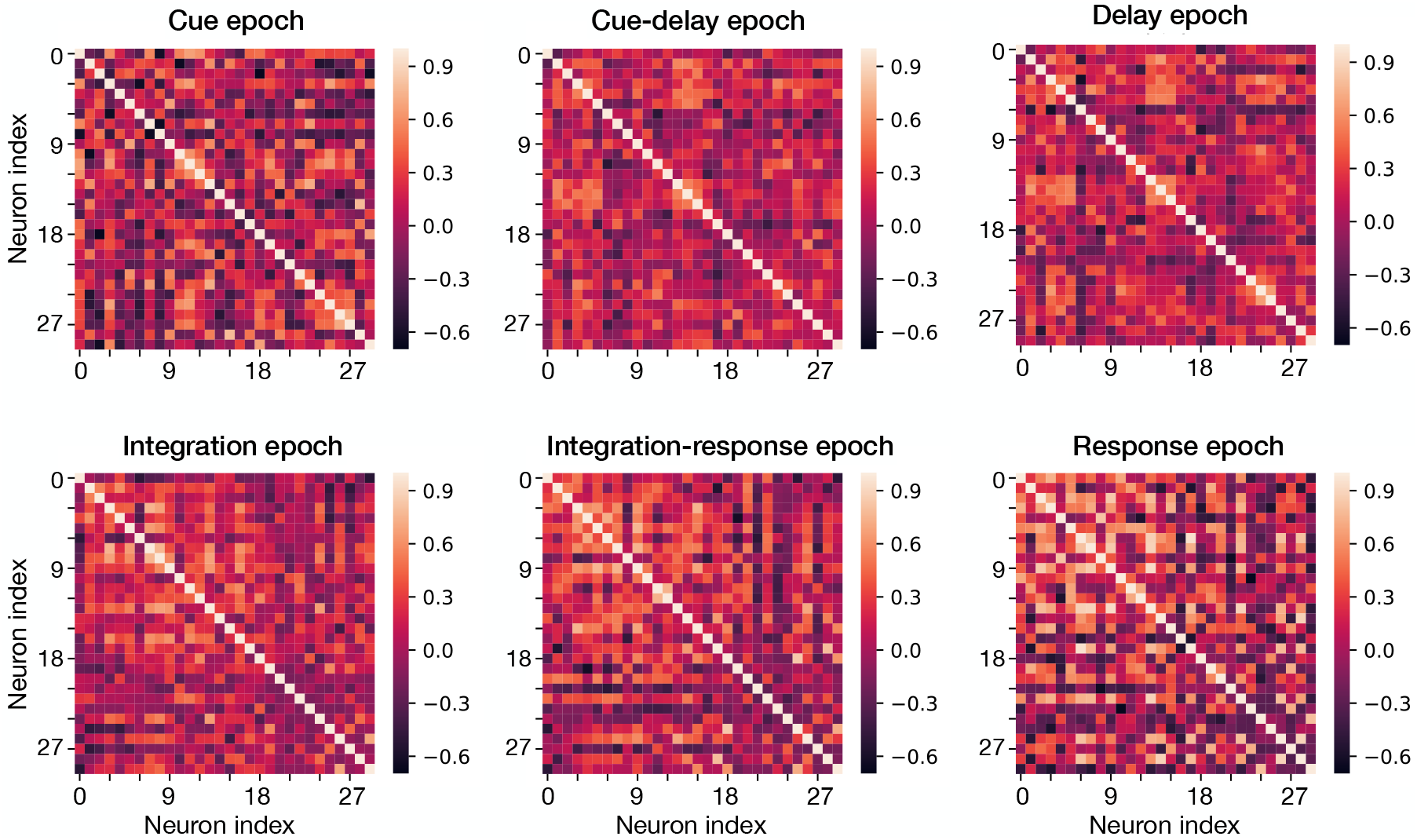
The correlation matrices of population activity in six task-specific periods. For clear visualization and comparison, we plotted 30 units with the higher mean activity across six epochs.

**Figure S6:**
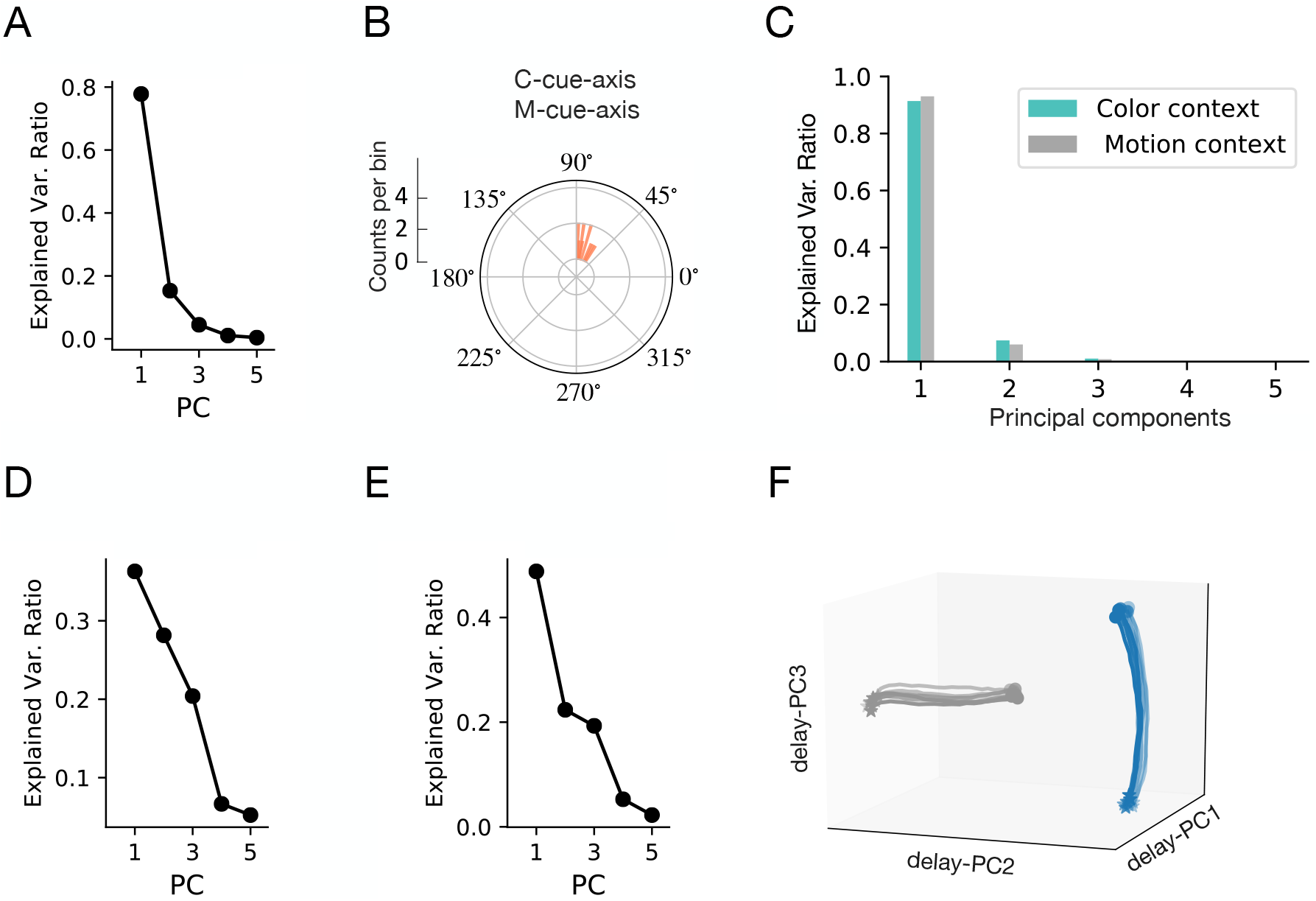
The impact of weight perturbation on neural representation in the subspace. The neural trajectory for four different SVs in a three-dimensional subspace, which is spanned by the first three PCs (delay-PC1,delay-PC2,delay-PC3). **(A)** Weight perturbation was operated during the complete task period, namely, global perturbation. The first/third-column panels indicate non-diagonal weight perturbation, and the second/fourth-column panels indicate diagonal weight. **(B)** Perturbation was operated only locally during the delay epoch.

**Figure S7:**
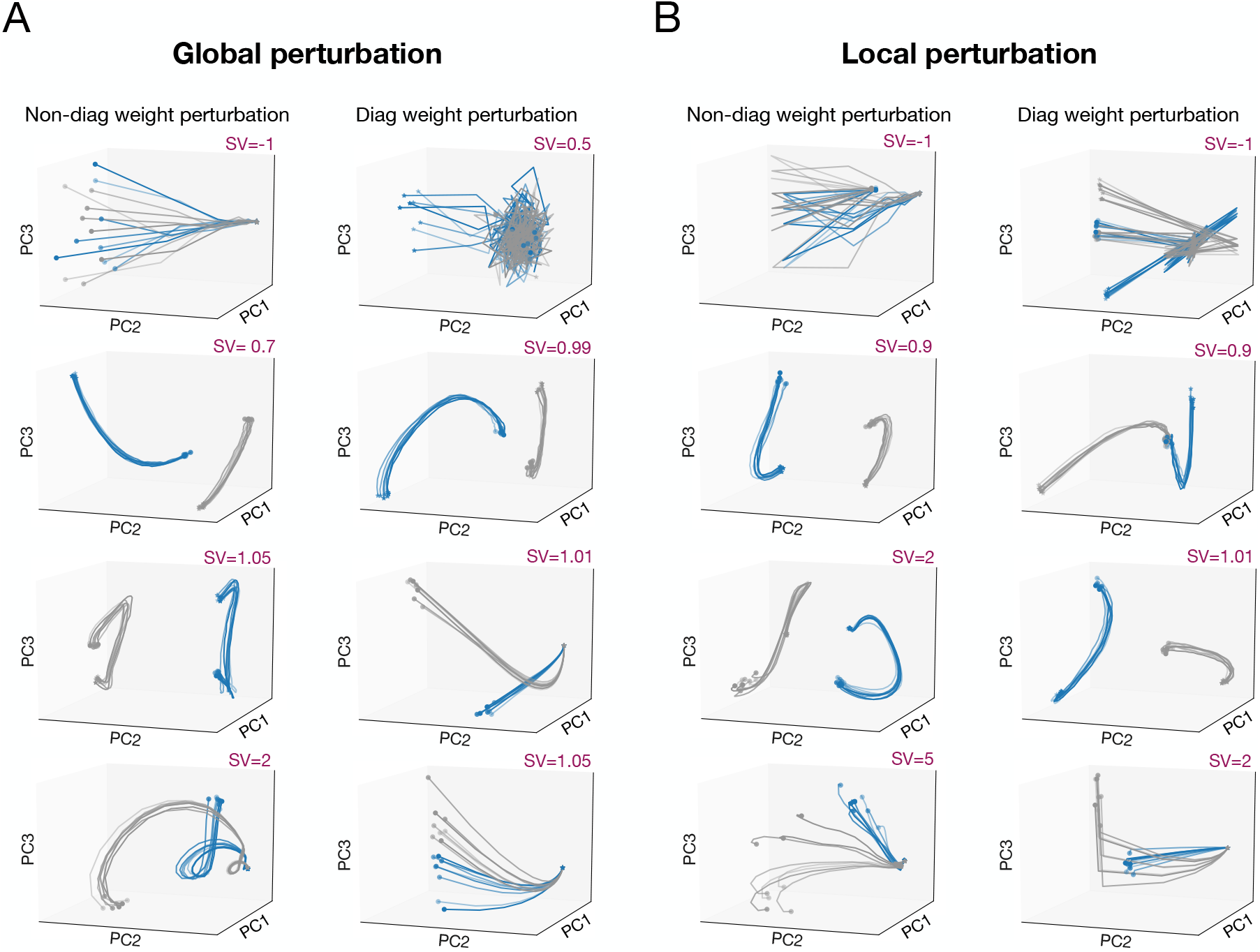
The impact of weight perturbation on neural representation in the subspace. The neural trajectory for four different SVs in a three-dimensional subspace, which is spanned by the first three PCs (delay-PC1,delay-PC2,delay-PC3). (A) Weight perturbation was operated during the complete task period, namely, global perturbation. The first/third-column panels indicate non-diagonal weight perturbation, and the second/fourth-column panels indicate diagonal weight. (B) Perturbation was operated only locally during the delay epoch.

**Figure S8:**
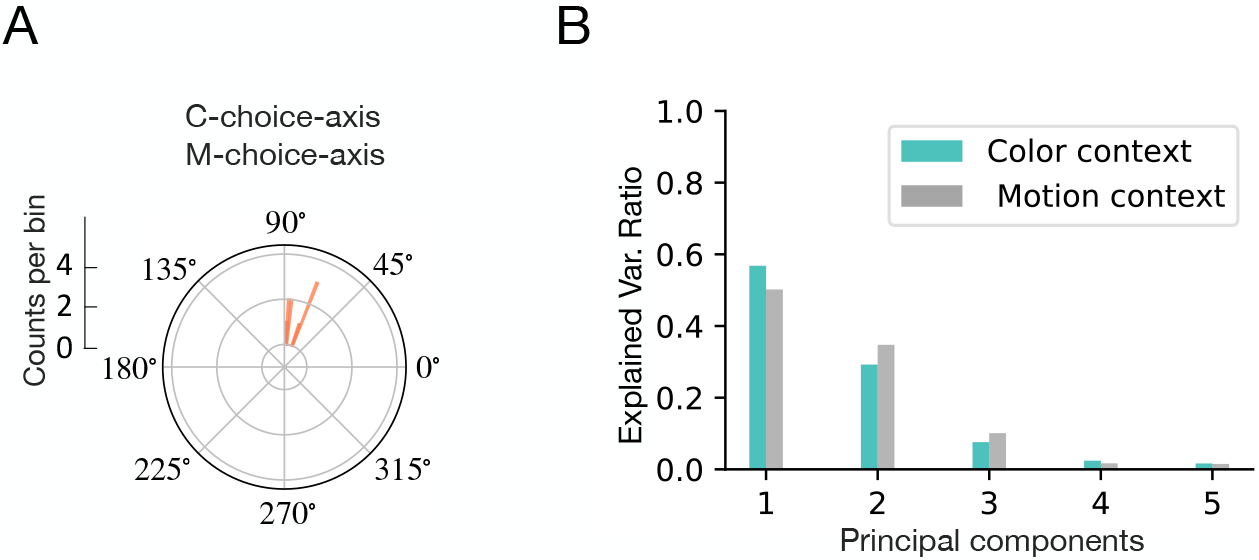
**(A)** Polar histogram of the angle between the `M-choice-axis` and `C-choice-axis`. **(B)** Ratio of explained variance of the first five PCs of subspace during the sensory stimulus epoch for the color and motion contexts, respectively.

**Figure S9:**
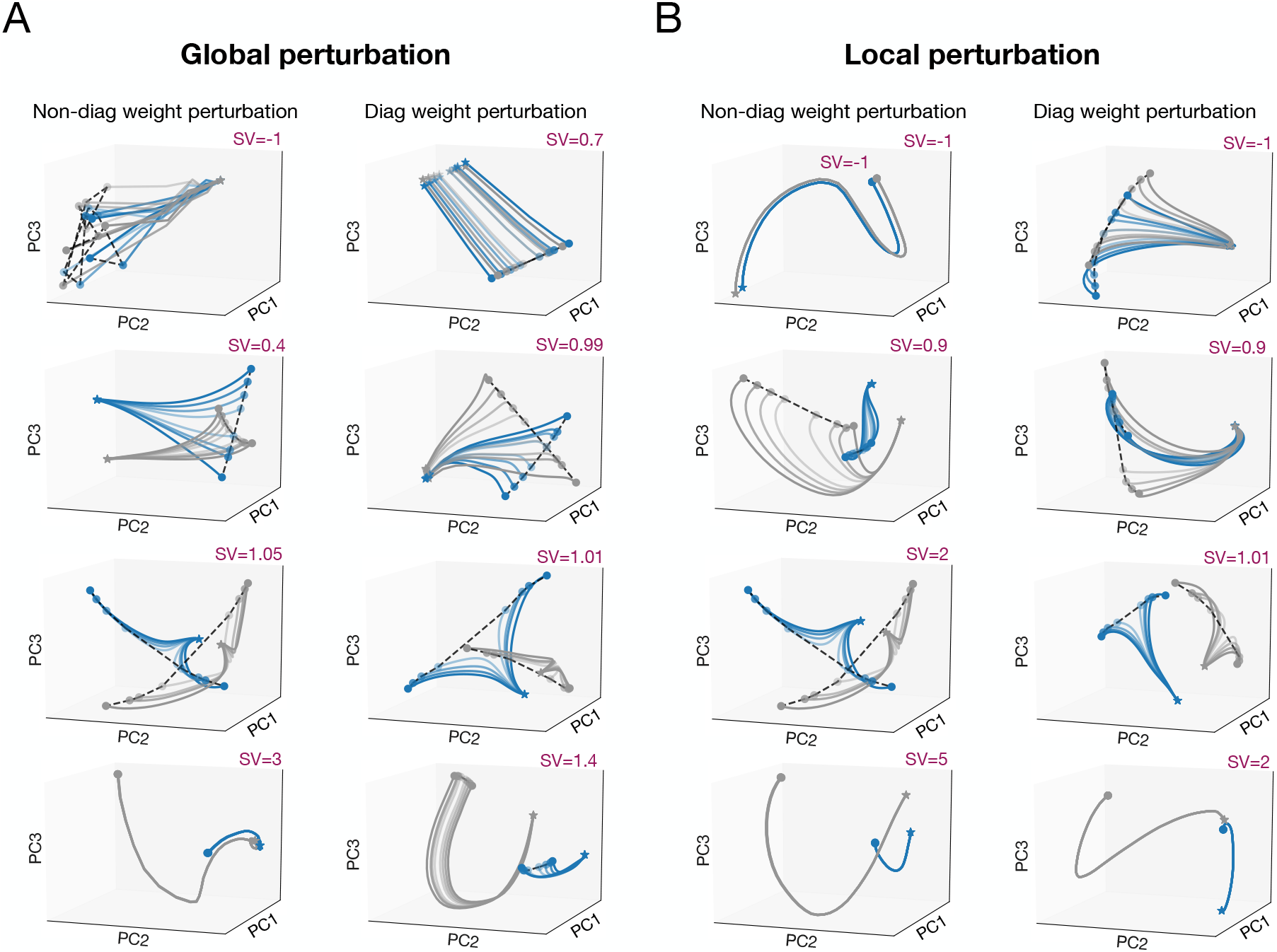
The neural trajectories under recurrent weight perturbation with different SV values. **(A)** Global recurrent weight perturbation was operated during the whole task period. The *left*column indicates non-diagonal weight perturbation, and the *right* column indicates diagonal weight. **(B)** Local perturbation was applied only locally in time during the delay epoch.

**Figure S10:**
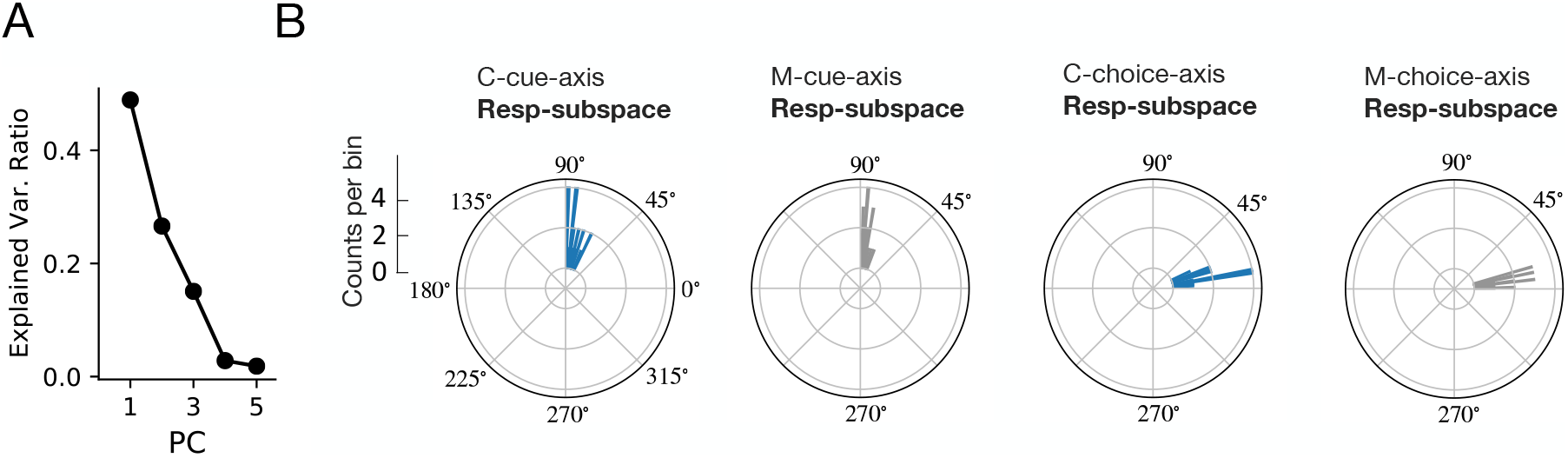
**(A)** Ratio of explained variance of the first five PCs of subspace during the response epoch. **(B)** The angle between four task-related axes and Resp-subspace.

**Figure S11:**
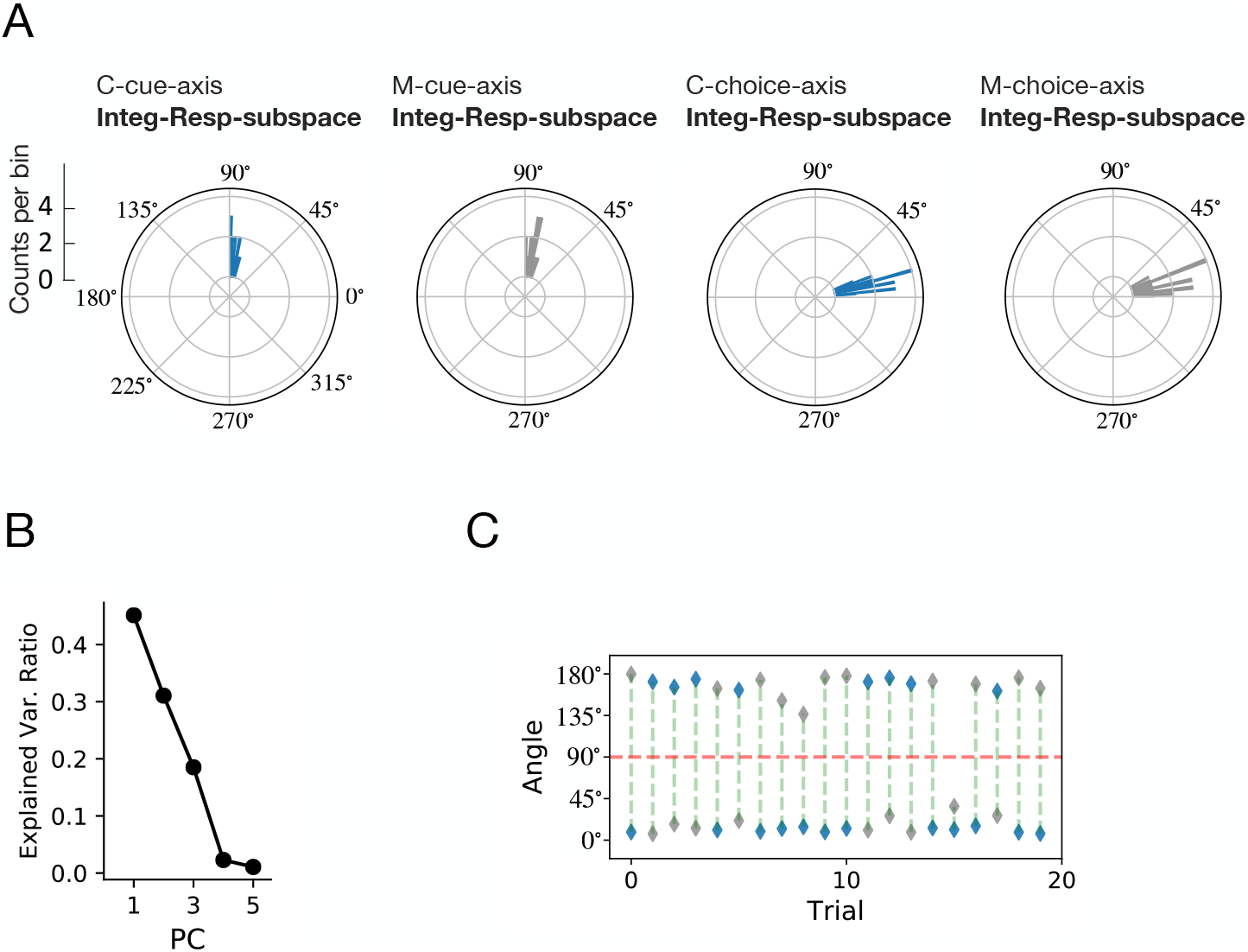
**(A)** The angle between four task-related axes and Integ-Resp-subspace. **(B)** Ratio of explained variance of the first five PCs of subspace. **(C)** The angle between the `C-cue-axis` (blue) or `M-cue-axis` (grey) and integ-resp-PC1 in 20 trials.

